# Microevolution toward loss of photosynthesis: Mutations promoting dark-heterotrophic growth and suppressing photosynthetic growth in cyanobacteria

**DOI:** 10.1101/2024.04.08.588626

**Authors:** Shintaro Hida, Marie Nishio, Kazuma Uesaka, Mari Banba, Nobuyuki Takatani, Shinichi Takaichi, Haruki Yamamoto, Kunio Ihara, Yuichi Fujita

**Affiliations:** Graduate School of Bioagricultural Sciences, Nagoya University, Nagoya, 464-8601, Japan; Center for Gene Research, Nagoya University, Nagoya, 464-8602, Japan; Department of Molecular Microbiology, Faculty of Life Sciences, Tokyo University of Agriculture, Tokyo, 156-8502, Japan

## Abstract

The prevalence of parasitic plants suggests frequent evolution of photosynthetic capacity loss in the natural environment. However, no studies have observed such evolutionary events as a loss of photosynthetic capacity. Herein, we report mutations that lead to a loss or decrease in photosynthetic growth capacity of dark-adapted variants of the cyanobacterium *Leptolyngbya boryana*, which can grow heterotrophically even in the dark. We isolated 28 dark-adapted variants through long-term cultivation (7–49 months) under dark-heterotrophic conditions. All variants showed significantly faster dark-heterotrophic growth than the parental strains, accompanied by the loss of photosynthetic growth capacity in 15 variants. Genome resequencing of the variants revealed that 19 of the 28 variants carried various mutations in a common single gene (*LBDG_21500*) encoding a protein phosphatase 2C (PP2C) RsbU that is involved in the partner switching system (PSS). Phenotypic and transcriptomic analyses of a *LBDG_21500*-knockout mutant suggested that the PSS, including LBDG_21500, is involved in the global transcriptional regulation of various genes under both photoautotrophic and dark-heterotrophic conditions. We propose the renaming of *LBDG_21500* to *phsP* (phototrophic–heterotrophic switching phosphatase). Our results imply that mutations in the global transcriptional regulatory system serve as the first evolutionary step leading to the loss of photosynthetic capacity.

**Importance:** Photosynthetic organisms that grow using minimal resources: light, water, and CO_2_, support most heterotrophic organisms as producers on the Earth. When photosynthetic organisms thrive over long generations under environments where organic compounds are readily available, they may lose the photosynthetic capacity because of the relief of selective pressure to maintain photosynthesis. The prevalence of parasitic plants in the natural environment supports this idea. However, there have been no actual observations of evolutionary processes leading to a loss of photosynthetic growth capacity. The significance of our research is in observing microevolution of a cyanobacterium through a long-term cultivating under dark heterotrophic conditions. In particular, the high frequency of mutations to a gene involved in the global transcriptional regulatory system suggests that such mutations in regulatory systems are regarded as an example of the initial evolutionary processes toward complete loss of photosynthesis.

## Introduction

Photosynthesis is the biochemical process underlying photoautotrophic growth, which uses light as energy and CO_2_ as a carbon source. Photoautotrophic organisms such as cyanobacteria, algae, and plants have the ability to grow with minimal resources such as light (energy), water (reducing power), and air (carbon source) and support most heterotrophic organisms as producers. However, in environments where organic compounds such as glucose are readily available, photoautotrophic organisms with chemoheterotrophic activity can use glucose as an energy and carbon sources; therefore, photosynthesis is not necessarily required as long as glucose is supplied. Under such an environment, selective pressure to maintain photosynthesis is relieved, and organisms may lose the photosynthetic capacity to survive. In fact, there are many non-photosynthetic species that have lost their photosynthetic capacity in the lineages of photosynthetic organisms; for example, parasitic plants such as beachdrops (*Epifagus virginiana*, de Pamphilis and Palmer, 1990; Wolfe et al. 1992); various members of the broomrape family (Wicke et al. 2013); the genus *Rafflesia* (e.g., *Raflesia lagascae*, Molina, 2014); mycoheterotrophs such as *Aneura mirabilis* (Wickett et al. 2008), *Sciaphila thaidanica* (Peterson et al. 2018), and *Monotropastrum humile* (Graham et al. 2017); the marine cyanobacterium UCYN-A in a symbiotic relationship with coccolithophytes (Tripp et al. 2010); *Nostoc azolla* in a symbiotic relationship with water ferns (Ran et al. 2010); endosymbiotic cyanobacteria (spheroid bodies) of diatoms (Nakayama et al. 2014); nonphotosynthetic *Cryptomonas* (Tanifuji et al. 2020); and the parasitic protozoan *Plasmodium falciparum* (Wilson et al. 1996). In all these cases, continued dependence on the supply of carbon from the host organism through a long-term symbiotic or parasitic relationship is assumed to have resulted in the relaxation of functional constraints on the genes involving in photosynthesis, allowing mutations to accumulate, leading to pseudogenization and loss of the responsible genes, ultimately leading to the loss of photosynthetic capacity. Plastid genome comparisons in parasitic plants (Wicke et al. 2013) and *Cryptomonas* algae (Tanifuji et al. 2020), and the substantial genome streamlining observed in cyanobacteria (Tripp et al. 2010) support such an evolutionary process. However, there have been no actual observations of evolutionary processes leading to a loss of photosynthetic growth capacity.

The partner switching system (PSS), which is widely distributed in prokaryotes, is a transcriptional regulatory system that controls various cellular processes such as the stress response, sporulation, biofilm formation, and pathogenicity (Yang et al. 1996; Bouillet et al. 2018). The PSS consists of four components: a sigma factor, an anti-sigma factor (with a kinase module), an anti-sigma antagonist, and a protein phosphatase 2C (PP2C). In the absence of specific signals for the PPS, the anti-sigma antagonist does not bind to the anti-sigma factor in its phosphorylated state, the anti-sigma factor binds to the sigma factor, and no sigma factor-dependent transcription of the regulon genes occurs. Once a specific signal is sensed, the phosphatase dephosphorylates the anti-sigma antagonist and then the dephosphorylated anti-sigma antagonist binds to the anti-sigma factor, resulting in the release of the sigma factor from the anti-sigma factor, which recruits RNA polymerase to initiate transcription of the regulon genes.

Cyanobacteria are photoautotrophic prokaryotes that perform oxygenic photosynthesis in a similar manner to plants, and most of them are obligate phototrophs (Rippka et al. 1979; Zhang et al. 1998). Some cyanobacteria exhibit chemoheterotrophic growth capacity in the dark using sugars such as glucose (White and Shilo 1975). The cyanobacterium *Leptolyngbya boryana* IAM M-101 can grow heterotrophically even in the dark in the presence of glucose (Fujita et al. 1992; Fujita et al. 1993), and the doubling time of the wild-type (WT) is substantially long (approximately 103 h). A spontaneous variant, known as dg5, which has a shorter doubling time (approximately 68 h) in dark-heterotrophic growth conditions, was isolated through short-term dark-heterotrophic cultivation (40 days; Fujita et al. 1996). Because dg5 maintains normal photosynthetic growth capacity, it provides a novel platform for investigating the physiological aspects of cyanobacteria in the dark, which is exemplified by the identification of *chlB* encoding a subunit of dark-operative protochlorophyllide oxidoreductase (DPOR) (Fujita et al. 1996) and the analysis of the chlorophyll deficiency process (etiolation) in the dark and subsequent light-dependent greening upon light irradiation in a mutant lacking DPOR (Kada et al. 2003). A frameshift in the gene *cytM* encoding cytochrome *c*_M_ was identified as the mutation responsible for the dg5 phenotype by determining the genomes of dg5 and the WT (Hiraide et al. 2015).

In this study, we isolated 28 spontaneous variants from *L. boryana* dg5 and the WT through long-term cultivation under dark-heterotrophic conditions (7–49 months). These variants commonly exhibited the phenotype of faster heterotrophic growth in the dark and slower photosynthetic growth in the light compared to the parental strains, and 15 of them lost photosynthetic growth capacity in the light. Genome resequencing of the variants revealed that many of them (19 of 28 variants) carried various mutations in a gene encoding a PP2C-type phosphatase, which was assumed to be a component of the PSS. Phenotype analysis of a targeted mutant in which the PP2C-type phosphatase gene was knocked out and transcriptomic analysis of the mutant suggested that the PP2C-type phosphatase is involved in the global regulation of genes in the two growth modes—photoautotrophic and dark-chemoheterotrophic—in response to the respective growth conditions in *L. boryana*. A PSS that includes the PP2C-type phosphatase could play a pivotal role in global transcriptional regulation to optimize photoautotrophic and chemoheterotrophic growth. The accumulation of various mutations in the phosphatase gene in dark-adapted variants implies that mutations in the regulatory system eventually lead to the loss of photosynthetic growth capacity, which could be regarded as an example of the initial evolutionary processes toward complete loss of photosynthesis.

## Results

### Isolation of dark-adapted mutants through long-term heterotrophic cultivation

To examine how mutations accumulate during long-term culture under dark-heterotrophic conditions in the cyanobacterium *L. boryana*, 28 colonies (6 from dg5, v1–v6; 22 from the WT, dg201–dg229) were isolated on initial agar plates under dark-heterotrophic conditions (Fig. S1A–C) and were cultivated under dark-heterotrophic conditions for a long period via inoculation every 10–14 days onto new agar plates. Variants v1–v6 from dg5 were maintained in dark-heterotrophic conditions for 49 months, and 16 (dg201–dg223) and 6 (dg224–dg229) variants from the WT were maintained for 15 or 7 months, respectively (Fig. S1D). Then, these dark-adapted variants were examined for photoautotrophic growth in the light and heterotrophic growth in the dark (Fig. 1). All variants exhibited significantly faster heterotrophic growth in the dark than the parental strains. However, under photoautotrophic conditions, many variants showed decreased growth, and some (15 variants) did not grow at all.

**Figure 1.**
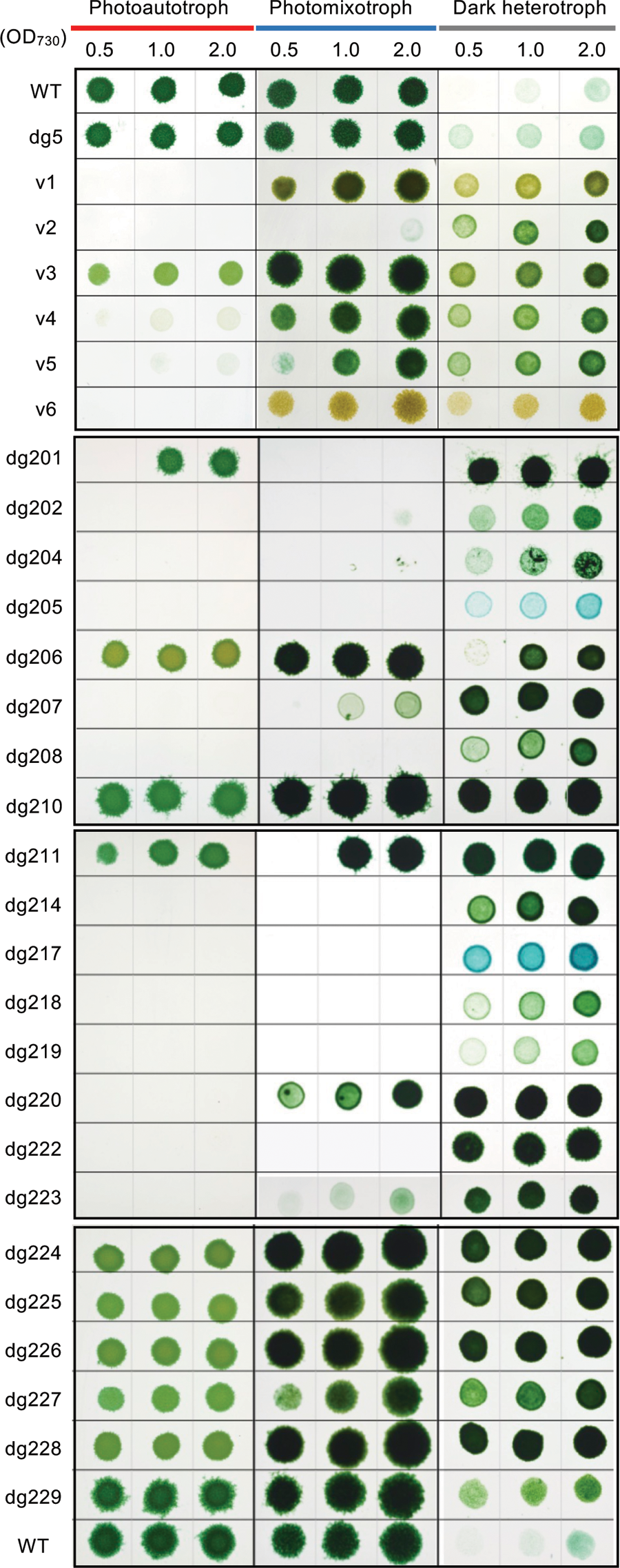
Growth comparison of dark-adapted variants v1–v6 and dg201–229 under photoautotrophic, photomixotrophic and chemoheterotrophic conditions. Cells of variants, wild type (WT) and dg5 were suspended in liquid BG-11 media to adjust the tubidity at OD_730_ to be 0.5, 1.0 and 2.0. Aliquots (10 µL each) were spotted on agar plates under photoautotrophic (BG-11; 20 µmol m^−2^ s^−1^), photomixotrophic (BG-11 containing 30 mM glucose; 20 µmol m^−2^ s^−1^), and heterotrophic (BG-11 containing 30 mM glucose; dark) conditions. All agar plates were incubated for 7 days.

The cellular absorption spectra and chlorophyll (Chl) contents of the dark-adapted variants grown heterotrophically in the dark were measured (Figs. S2 and S3). Overall, a decrease in phycobilin and Chl contents was a common characteristic of all variants, which was reflected in the unique color of the cultures, such as khaki in v1 and v6 and blue in dg205 and dg217 (Fig. 1). In particular, v6 showed the lowest peaks at approximately 630 and 680 nm, indicating the lowest contents of phycobilin and Chl among all variants (Fig. S2A, B). A significant decrease in carotenoid content was observed in three of the variants (dg205, dg211, and dg217) (Fig. S3A, B).

### Genome resequencing of the dark-adapted variants

Genome resequencing was performed to identify the mutations accumulated in the 28 dark-adapted variants. A total of 22 mutations were found in the 6 variants (v1 to v6) compared with the genome of the parental strain dg5 (Fig. 2A, Table S1). Additionally, 68 mutations were found in the 22 dark-adapted variants (dg201 to dg229) derived from the WT (Fig. 2B, Table S2). Excluding duplicate mutations in different variants, 63 independent mutations were found (Table S2). Interestingly, 19 of the 28 variants had mutations in the common gene *LBDG_21500* (dg5) / *LBWT_21500* (WT) (Fig. 2AB, Table S1–S3). Because the nucleotide sequences of *LBDG_21500* and *LBWT_21500* are identical, they are referred to *LBDG_21500* uniformly hereafter. These mutations were all different except for dg206 and dg217, which were identical (W364>Stop), resulting in 18 different types of mutations. Of these 18 mutations, 13 appear to cause loss of function in *LBDG_21500* (five Tn insertions, two deletions of relatively long fragments (1,197 bp and 993 bp), four frameshift mutations, and two nonsense mutations; Fig. 2C). The other five were single nucleotide substitutions for nonsynonymous amino acid substitutions (Fig. 2C). Among the 19 variants with mutations in *LBDG_21500*, nine variants showed no photosynthetic growth, and 10 variants showed severely or significantly decreased growth under photosynthetic conditions. The degree of growth retardation did not necessarily correlate with the type of mutation in *LBDG_21500*, suggesting that not only mutations in *LBDG_21500* but also other mutations combine to affect the growth characteristics (see Discussion). LBDG_21500 is annotated as “serine phosphatase RsbU, regulator of sigma subunit” in the database and corresponds to RsbU (regulator of <u>s</u>igma <u>B</u>) in *Bacillus subtilis* (Voelker et al. 1995). RsbU, a PP2C-type serine phosphatase, plays a role as the phosphatase unit catalyzing the dephosphorylation of the anti-sigma antagonist (RsbV) in the signal transduction PSS involved in activating the sigma factor (sigma B) in response to environmental stress in *B. subtilis* (Yang et al. 1996). Interestingly, most mutations in the gene LBDG_21500 occur in the C-terminal phosphatase domain (Fig. 2C), strongly suggesting that loss of function or decreased activity of the C-terminal phosphatase domain of the LBDG_21500 protein leads to slower photosynthetic growth in the light and faster heterotrophic growth in the dark compared to the parental strains. Additionally, mutations in *LBDG_21500* occurred in both parental strains, dg5 and the WT, indicating that such mutations are selected at a high frequency under dark-heterotrophic conditions regardless of the genetic background of *cytM* (the gene responsible for the dg5 phenotype; Hiraide et al. 2015).

**Figure 2.**
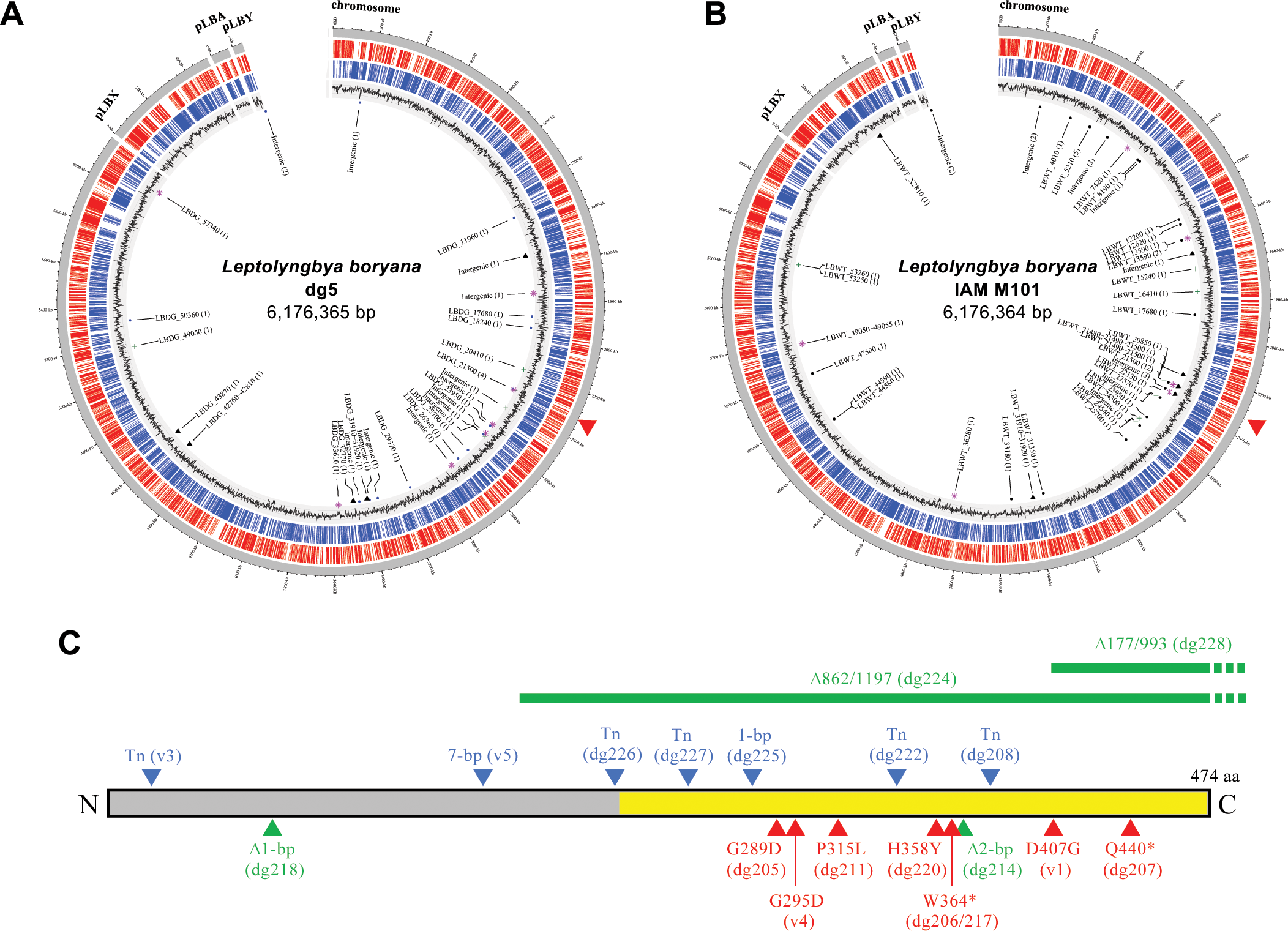
Circos plots of the genomes of six variants from dg5 (A) and 22 variants from the WT (B). ORFs of clockwise and counterclockwise directions are shown by red and blue lines, respectively. GC skew is shown in the third circles. Mutation types are shown indicated as follows: blue dot, SNP; green +, insertion; triangles, deletion; pink *, transposon insertion. The genes with the mutations are shown as locus tags. The numbers in the parentheses indicate the number of mutations occurring in the same genes in the v1–v6 (**A**) and dg201–229 (**B**) variants. The locus of *LBDG/LBWT_21500* is shown by the red triangles. (**C**) Distribution of mutations in the coding region of *LBDG_21500*. The C-terminal half corresponding to the phosphatase domain is shown in yellow. Insertions (Tn), deletions, and 1-bp substitutions are indicated by blue, green, and red triangles, respectively. Two long deletions covering the 3′-regions of *LBDG_21500* to the downstream chromosomal region are shown as green horizontal bars with dotted lines on the right side. The names of variants carrying the mutations are shown in the parentheses.

### Correlation between mutations and loss of photosynthetic growth capability

From the list of mutations in the 28 variants, the following two points are particularly noteworthy concerning the correlation with photosynthetic growth capacity. First, mutations in the genes *ccsA* (*LBDG_05210*) and *ccsB* (*LBDG_13590*) for cytochrome *c* biosynthesis (cytochrome *c* biosynthesis system II) (Simon and Hederstedt, 2011) appeared to correlate well with the loss of photosynthetic growth capacity. All nine variants (six with *ccsA* mutations and three with *ccsB* mutations) exhibited the common phenotype of no growth under photosynthetic conditions. Although all variants were accompanied by other mutations, *ccsA* and *ccsB* are probably essential for photosynthetic growth in *L. boryana*, which is consistent with a previous report that described that *ccsA* could not be disrupted in *Synechocystis* sp. PCC 6803 under photoautotrophic and photoheterotrophic conditions and that *ccsA* transcription is light-dependent (Hübschmann et al. 1997).

The other point is the essential role of carotenoids in photosynthetic growth. We performed pigment analysis for three variants that were presumed to have carotenoid deficiency based on the cellular absorption spectra (dg205, dg211, and dg217; Fig. 1, Fig. S3A, B). Pigments of the three variants were analyzed using high-performance liquid chromatography (HPLC) (Fig. 3). The WT of *L. boryana* contains β-carotene, zeaxanthin, ketomyxol glycoside (KMG), and echinenone as carotenoids (Fig. 3A). None of these carotenoids were detected in dg205 (Fig. 3B). Furthermore, no phytoene, the common precursor of all carotenoids, was detected in dg205 (Fig. 3B, E), indicating that dg205 contains no carotenoids. The HPLC profiles of dg211 showed accumulation of two *cis*-lycopenes and ζ-carotene instead of normal carotenoids (Fig. 3C, F), suggesting a deficiency in carotene isomerase (CrtH) activity. In dg217, only a small amount of phytoene and only a trace amount of phytofluene were detected (Fig. 3D, G), suggesting a significant decrease in the activities of phytoene synthase (CrtB) and phytoene desaturase (CrtP). Considering the mutations detected in genome resequencing, the loss of phytoene in dg205 agrees well with the loss of phytoene synthase activity due to a single adenine insertion in the coding region (frameshift) of *crtB* (*LBDG_53260*, Tables S2, S3). The abnormal accumulation of carotenoid intermediates in dg211 appears to be consistent with the loss of CrtH activity (carotene isomerase) (Fig. 3E; Masamoto et al. 2001), which is caused by a Tn insertion in the *crtH* coding region (*LBDG_36280*, Tables S2, S3). The pigment composition of dg217 suggests a significant decrease in CrtP and CrtB activities, although genome resequencing showed only a single nucleotide insertion in the latter half of the *crtP* coding region (*LBDG_53250*, Tables S2, S3). The frameshift in *crtP* occurs after Lys294 in the total length of 471 residues, and a small amount of residual activity by the truncated CrtP may only convert a small amount of phytoene to phytofluene, although the WT CrtP converts phytoene to ζ-carotene via phytofluene (Fig. 3E). Although, no mutation was detected in *crtB*, the low amount of phytoene (Fig. 3D) suggested that CrtB activity was also greatly decreased. The reason for this is currently unknown; however, the two genes *crtP* and *crtB* form an operon that is highly conserved in cyanobacteria. This implies that CrtP and CrtB are stabilized by their interaction with each other and that the truncation of CrtP results in a substantial decrease in the stability and/or activity of CrtB.

**Figure 3.**
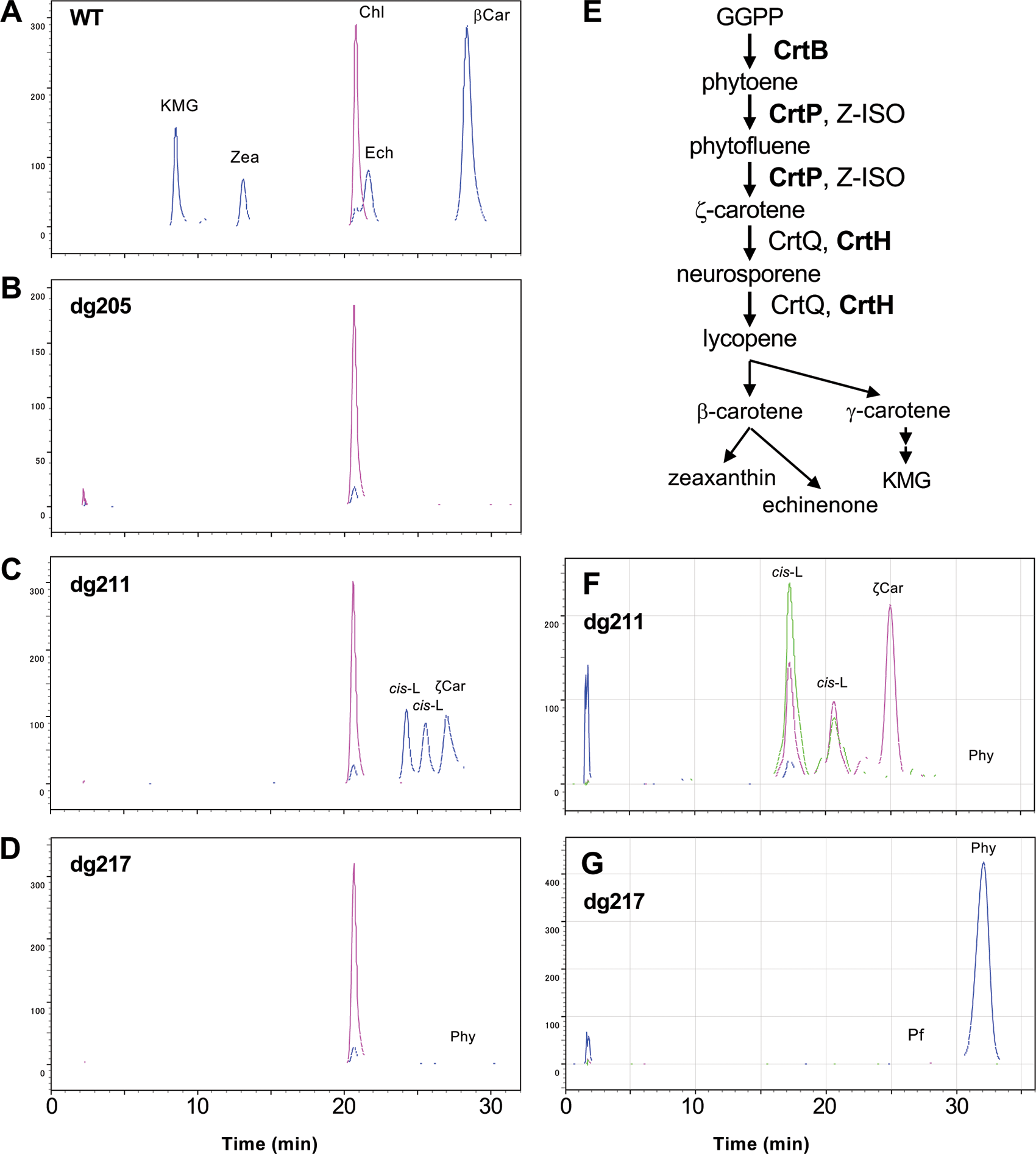
Pigment analysis of the dark-adapted variants, dg205 (**B**), dg211 (**C, F**), and dg217 (**D, G**), showing deficiency of carotenoids and WT (**A**). (**E**) Carotenoid biosynthesis pathway with the enzyme names. Total pigments were detected by absorbance at 618 nm (pink) and 470 nm (blue) in panels **A-D** using a pigment separation system. Only carotenes were enriched and detected at 285 nm (blue), 400 nm (pink), and 450 nm (green) in panels **F** and **G** using the carotene separation system. Chl, chlorophyll *a*; KMG, ketomyxol glycoside; Zea, zeaxanthin; βCar, β-carotene; Ech, echinonen; *cis*-L, *cis*-lycopene; ζCar, *d*ζ-carotene; Phy, phytoene; and Pf, phytofluene (see *Pigment analysis* in Experimental Procedures).

Interestingly, mutations in the *ccsA*, *ccsB*, *crtB, crtH*, and *crtP* genes were accompanied by mutations in *LBDG_21500* at a high frequency. Four of the six variants with mutations in *ccsA*, and two of the three variants with mutations in *ccsB* were accompanied by mutations in *LBDG_21500*. All three variants with mutations in *crt* were accompanied by mutations in *LBDG_21500*.

### Growth of a targeted LBDG_21500 mutant

To examine the effect of loss of function of LBDG_21500, *LBDG_21500*-knock out mutants (Δ21500/WT; Δ21500/dg5) were isolated (Fig. S4A, B) from the WT and dg5, and their growth was compared with that of the parental strains (Fig. 4). Both mutants showed commonly enhanced dark-heterotrophic growth accompanied by decreased photosynthetic growth. The changes in the Chl content in liquid culture (Δ21500/dg5) were examined (Fig. 4B). In Δ21500, the Chl content remained at a lower level than in dg5 under light conditions (Fig. 4B, b). This trend was reversed in the dark, with higher Chl content in Δ21500/dg5 than in dg5 (Fig. 4B, f). This difference in Chl content was also reflected in the cellular absorption spectra (see the 680 nm peak in Fig. 5A). In other words, dg5 dynamically changed the Chl content in response to growth conditions, whereas Δ21500 was less responsive to environmental changes and the Chl content did not change significantly under either condition.

**Figure 4.**
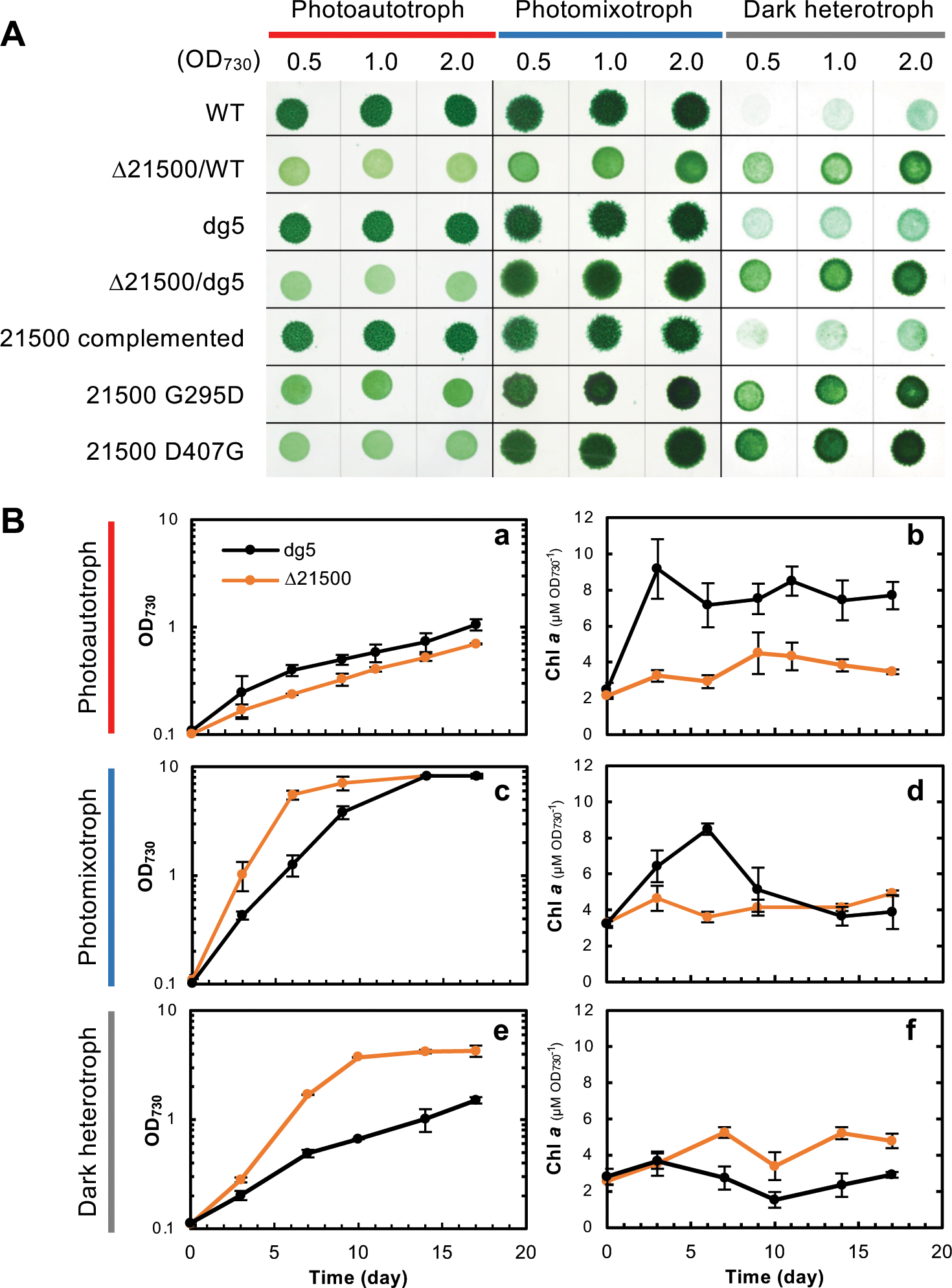
**(A)** Growth comparison of the targeted mutants of LBDG_21500 (Δ21500) derived from WT and dg5 with mutants carrying LBDG_21500 variants G295D and D407G under photoautotrophic, photomixotrophic, and dark-heterotrophic conditions on agar plates. (**B)** Growth (a, c, and e) and chlorophyll contents (b, d, and f) were compared in liquid cultures for Δ21500/dg5 (orange) and dg5 (black) under photoautotrophic (a and b), photomixotrophic (c and d), and dark-heterotrophic conditions (e and f).

**Figure 5.**
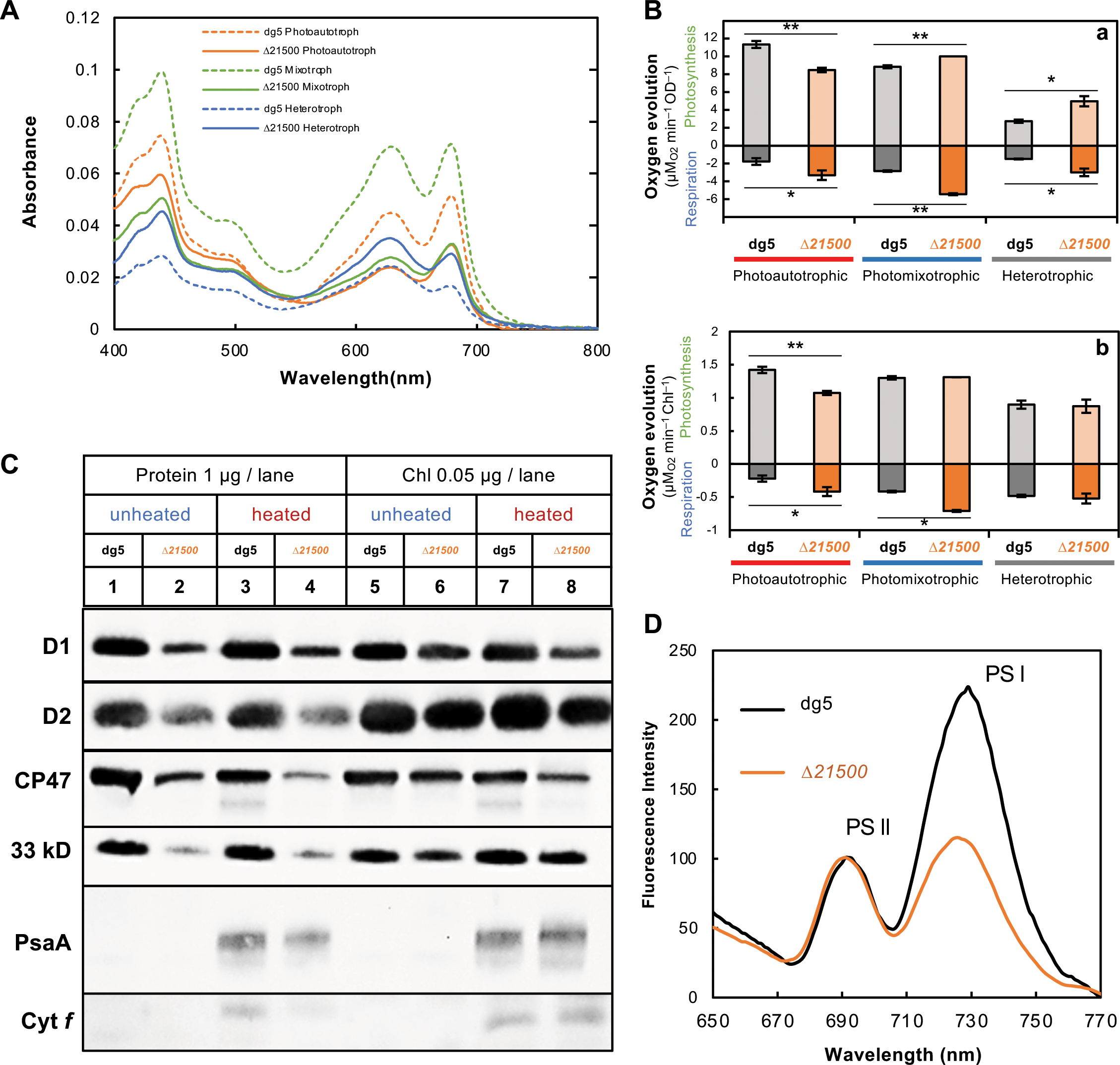
**(A)** Absorption spectra of dg5 and Δ21500 cells grown under photoautotrophic, photomixotrophic, and dark-heterotrophic conditions. (**B**) Activities of photosynthesis (oxygen evolution) and respiration (oxygen consumption) of targeted mutants of *LBDG_21500* (Δ21500) and dg5 grown under photoautotrophic, photomixotrophic, and dark-heterotrophic conditions. Activities are shown in OD_730_ (**a**) and chlorophyll (**b**) bases. (n=3). Bars represent standard deviations. Single and double asterisks indicate P < 0.05 and P < 0.001, respectively, between dg5 and Δ21500 (unpaired Student’s t-test followed by Benjamini–Hochberg multiple test corrections). (**C**) Western blot analysis of Δ*21500* (lanes 1, 3, 5 and 7) and dg5 (lanes 2, 4, 6 and 8) grown under photoautotrophic conditions. Samples were loaded onto each lane with a protein (lanes 1-4) or chlorophyll (lanes 5–8) base. Protein samples were unheated (lanes 1, 2, 5, and 6) or heated (lanes 3, 4, 7 and 8). Specific protein signals were detected using specific antisera against D1, D2, CP47, 33 kD (PsbO), PsaA, and cytochrome *f*. (**D**) Low temperature (77K) fluorescence spectra of Δ*21500* and dg5 with mutants grown under photoautotrophic conditions. Cells were adjusted to 0.5 µg_Chl_ ml^−1^. Fluorescence spectra were normalized to the peak of photosystem II (F690).

In addition to the knockout mutants of *LBDG_21500*, two *LBDG_21500* variants with two amino acid substitutions (G295D, D407G; the two mutations in v1 and v4) were introduced into Δ21500/dg5 to confirm functional complementation (Fig. S4C, D). A control strain carrying WT *LBDG_21500* at the same locus showed the WT phenotype, confirming full complementation by the WT copy. Meanwhile, the two amino acid substitution mutants (G295D and D407G) exhibited the same phenotype as the Δ21500 mutants (Fig. 4A), indicating that both D407G and G295D substitutions cause the same effect as the loss of function of the LBDG_21500 protein.

These results suggest that the LBDG_21500 protein is involved in the regulation of photosynthetic growth in the light and heterotrophic growth in the dark. Given that the PSS consists of four components, including a PP2C-type serine phosphatase, LBDG_21500 could be a member of a PSS regulatory system that promote photosynthetic growth and suppress dark-heterotrophic growth in *L. boryana*. Of the 19 variants with mutations in *LBDG_21500*, only 11 exhibited the same phenotype as Δ21500, suggesting that mutations other than *LBDG_21500* in the other eight variants may have more negative effects on photosynthetic growth (Table S3, see Discussion).

### Analysis of photosystems in Δ21500

To investigate the causes of poor photosynthetic growth and better dark heterotrophic growth in Δ21500, activities of photosynthesis (oxygen production) and respiration (oxygen consumption) were compared with those of the parental strain dg5 (Fig. 5B). Because the Chl contents of Δ21500 and dg5 were different, the activities were compared per turbidity (optical density (OD), panel a) value and per Chl content (panel b). Under photoautotrophic conditions, the activity of photosynthesis of Δ21500 was significantly lower both per Chl content and per OD value than that of dg5, which was consistent with poor photosynthetic growth. Cells grown under dark heterotrophic conditions, where respiration is the primary process of energy production, showed higher respiration activity per OD value in Δ21500 than dg5, which is consistent with the faster heterotrophic growth of Δ21500 in the dark. However, it should be noted that the respiratory activity of Δ21500 was not significantly different when compared on a per Chl content basis because of the higher Chl content under dark-heterotrophic conditions in Δ21500.

Next, Western blot analysis of the membrane fractions of Δ21500 and dg5 cells grown under photosynthetic conditions was performed using specific antibodies against subunit proteins of the photosynthetic electron transfer system: D1, D2, CP47, and 33 kDa (for photosystem II); PsaA (for photosystem I); and cytochrome *f* (for cytochrome *b*_6_*f*) (Fig. 5C; Fig. S5). Again, because the Chl contents of Δ21500 and dg5 were not the same, the amount of sample loaded for SDS-PAGE was compared per protein (Fig. 5C, lanes 1–4; 1 µg/lane) and per Chl content (Fig. 5C, lanes 5–8; 0.05 µg_Chl_/lane). Additionally, heat-treated and non-heat-treated samples were also subjected to this analysis because some photosystem subunits change their behavior on SDS-PAGE upon heat treatment. The results showed that on a per-protein basis, all proteins examined were significantly lower in Δ21500. However, because the Chl content was decreased in Δ21500, the difference in protein contents between Δ21500 and the WT was not as pronounced on a per-Chl basis.

To obtain information on the ratio of the photosystems, we measured the low-temperature fluorescence spectra of Δ21500/dg5 and dg5 cells grown under photosynthetic conditions (Fig. 5D). The fluorescence spectra of cells with the same Chl content were normalized to the peak of photosystem II (F690). The fluorescence peak indicative of photosystem I (F730) in Δ21500 was approximately half of that in dg5, suggesting that photosystem I (relative to photosystem II) is significantly reduced in Δ21500/dg5 compared with dg5.

These results imply that the decrease in growth of Δ21500 under photosynthetic conditions was largely due to a relative decrease in photosystem I or a ratio imbalance between photosystems I and II. Additionally, the decrease in phycobiliproteins (Fig. 5A) was also presumed to contribute to poor growth in Δ21500 under photosynthetic conditions (see the RNA-seq analysis section).

### RNA-seq analysis of Δ21500 and dg5

To identify the gene set regulated by the signaling pathway involving LBDG_21500, dg5 and Δ21500/dg5 were grown under photosynthetic and dark-heterotrophic conditions, and transcriptome analysis (RNA-seq) for the four RNA samples (photosynthetically grown dg5 and Δ21500, and dark-heterotrophically grown dg5 and Δ21500) was performed in biological triplicate (Fig. 6A, Table S6). Clustering analysis of the 12 transcriptional profiles for a total of three replicates in the two conditions showed two major clusters under photosynthetic and dark-heterotrophic conditions. This indicates that the transcriptional profile changes significantly under light-photosynthetic and dark-heterotrophic conditions and that the effect of the loss of LBDG_21500 is smaller than this global change. Next, the two clades (light-photosynthetic and dark-heterotrophic) were further clustered by the presence or absence of LBDG_21500, dg5, and Δ21500. The results of the principal component analysis (PCA) were also consistent with the clustering results (Fig. S6). The transcript profiles changed significantly with growth mode (light-photosynthetic and dark-heterotrophic growth) and the presence or absence of LBDG_21500. This suggests that LBDG_21500 is partially responsible for global transcriptional regulation under both light-photosynthetic and dark-heterotrophic conditions.

**Figure 6.**
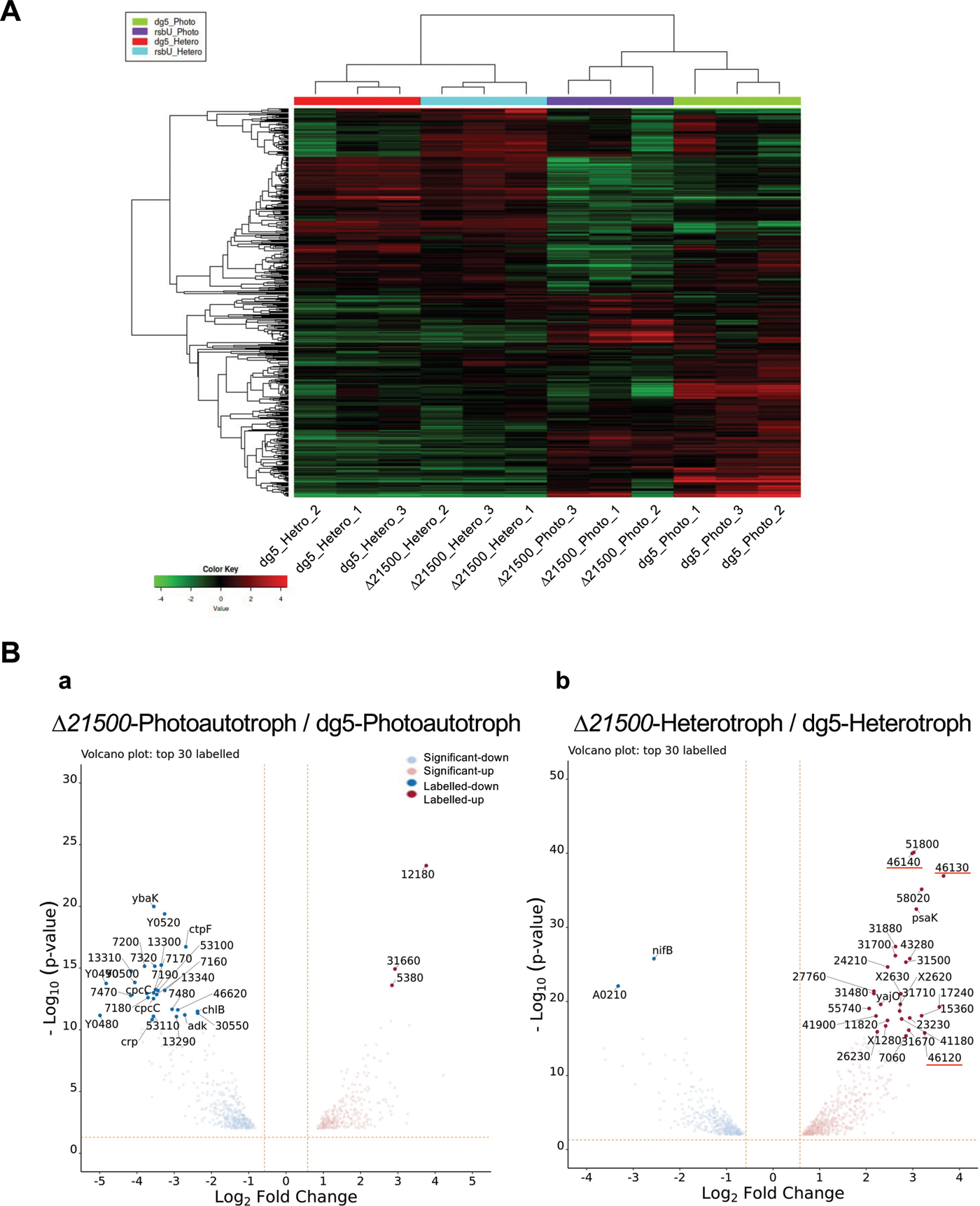
**(A)** Hierarchical clustered heatmap of the top 500 genes with large expression variability across twelve RNA seq datasets. Using the Log_2_ count per million (CPM) transformed expression count as input, the top 500 genes with large expression variation (SD) across all samples were analyzed. Hierarchical clustering was performed using an average linkage clustering method with correlation-based distance metrics. (**B**) Volcano plots of genes significantly differentially expressed by the knockout of *LBDG_21500* under light and dark conditions. Differentially expressed genes (DEGs) with higher and lower transcript levels in Δ*21500* are shown in red and blue, respectively, compared with dg5 under photoautotrophic (**a**) and with dark heterotrophic (**b**) conditions are shown in red and blue, respectively. Volcano plots were created based on the bases of the ratios of transcript levels (log_2_ fold change) and minus log_10_ transformed q-values of the genes with significantly differential expression at a significance level of q-value (FDR) < 0.01. The top 30 DEGs in the minus log_10_ transformed q-value were labeled; genes are shown with their gene names or the five digits after LBDG (locus tag). The dashed lines indicate the cut-off values for FDR (horizontal) and log_2_ fold-change (vertical).

Next, to clarify the target genes of transcriptional regulation involving LBDG_21500, comparative analysis of the transcript levels of each gene in Δ21500 and dg5 for light-photosynthetic and dark-heterotrophic conditions were performed (Fig. 6B, Figs. S7, S8, and S9). First, we compared the transcript levels of Δ21500 relative to those of dg5 under light-photosynthetic conditions and found that the transcript levels of 710 genes were significantly altered (Fig. 6Ba, Fig. S7A). Most (27 genes) of the top 30 genes that varied showed significantly decreased transcript levels (Fig. 6Ba, Table S7). When these 710 fluctuating genes were examined via gene set enrichment analysis (GSEA) (Fig. S9) in the Kyoto Encyclopedia of Genes and Genomes (KEGG) pathway. Genes for the KEGG functional categories for ribosome, photosynthesis antenna, photosynthesis, biosynthesis of secondary metabolites, oxidative phosphorylation were significantly enriched at a false discovery rate (FDR) level of <0.01 level (Fig. S9A). The decrease in the transcripts in these KEGG categories suggests that LBDG_21500 is involved in the transcriptional activation of genes involved in photosynthesis and the respiratory electron transport. Among these genes, genes encoding some phycobiliproteins (*cpcGBAC1C2*, LBDG_43870-43910 and *cpcC*, LBDG_26830, and *apcC*, LBDG_44410; Fig. 7) showed a significant decrease in transcript levels in Δ21500 grown under photosynthetic conditions. In the genome of *L. boryana*, there are four genetic loci for phycobiliproteins. *cpcBAC1C2* (Fig. 7B) and *cpcC* (Fig. 7A) showed significantly lower transcript levels in Δ21500 grown under photosynthetic conditions, and these two loci are corresponding to the two *cpc* operons with the highest transcript levels of dg5 under photosynthetic conditions. Therefore, the decrease in transcript levels of the *cpc* and *apc* genes appears to be consistent with the decrease in the absorption peak for phycobiliproteins in Δ21500 cells under photosynthetic conditions was observed (Fig. 5A), suggesting that the decreased transcript levels of the major *cpc* operons in Δ21500 is largely responsible for the reduced phycobilin levels.

**Figure 7.**
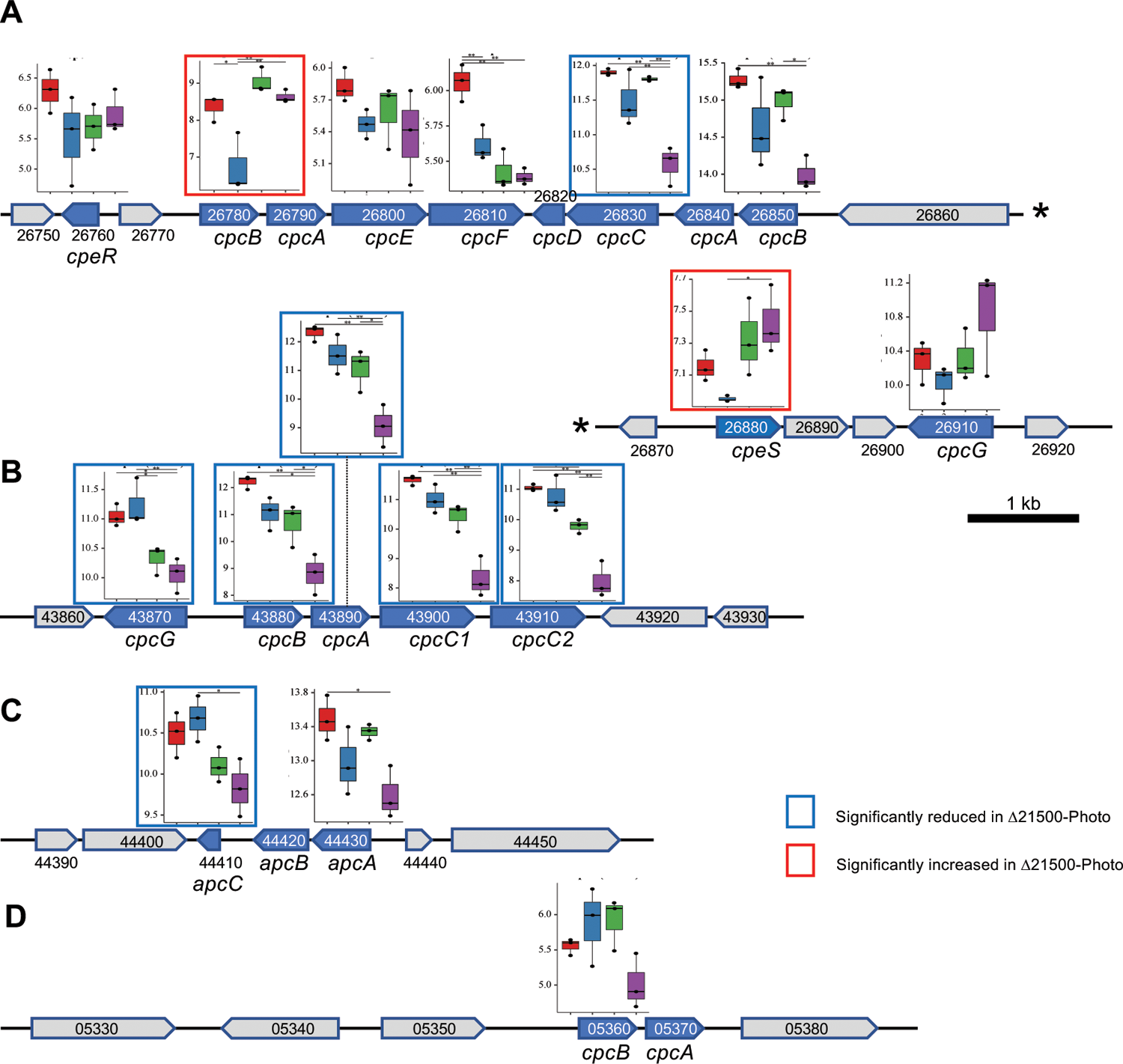
Four genetic loci (**A**–**D**) encoding phycobiliproteins in *L. boryana* with transcript profiles of Δ*21500* and dg5 grown under photoautotrophic and heterotrophic conditions. Genes encoding phycobiliproteins are shown in blue. The transcript profiles of the genes are shown just above the gene. Transcript profiles are shown in log_2_ expression levels, with log-transformed TPM-normalized expression values in four conditions (red, dg5/heterotrophic; blue, dg5/photoautotrophic; green Δ21500/heterotrophic; and violet, Δ21500/photoautotrophic). Each plot shows the mean of three replicates. Asterisks indicate significant differences between each pair at the 5% risk level (*) or 1% risk level (**) after Benjamini–Hochberg (BH) multiple testing correction for Student’s *t*-test. Statistical tests and box-and-whisker plots were performed using the FaDA shiny server (Danger, 2021). Profiles showing significant increases and decreases in Δ*21500* under photoautotrophic conditions (comparison between blue and violet boxes) are shown by blue and red squares, respectively.

**Figure 8.**
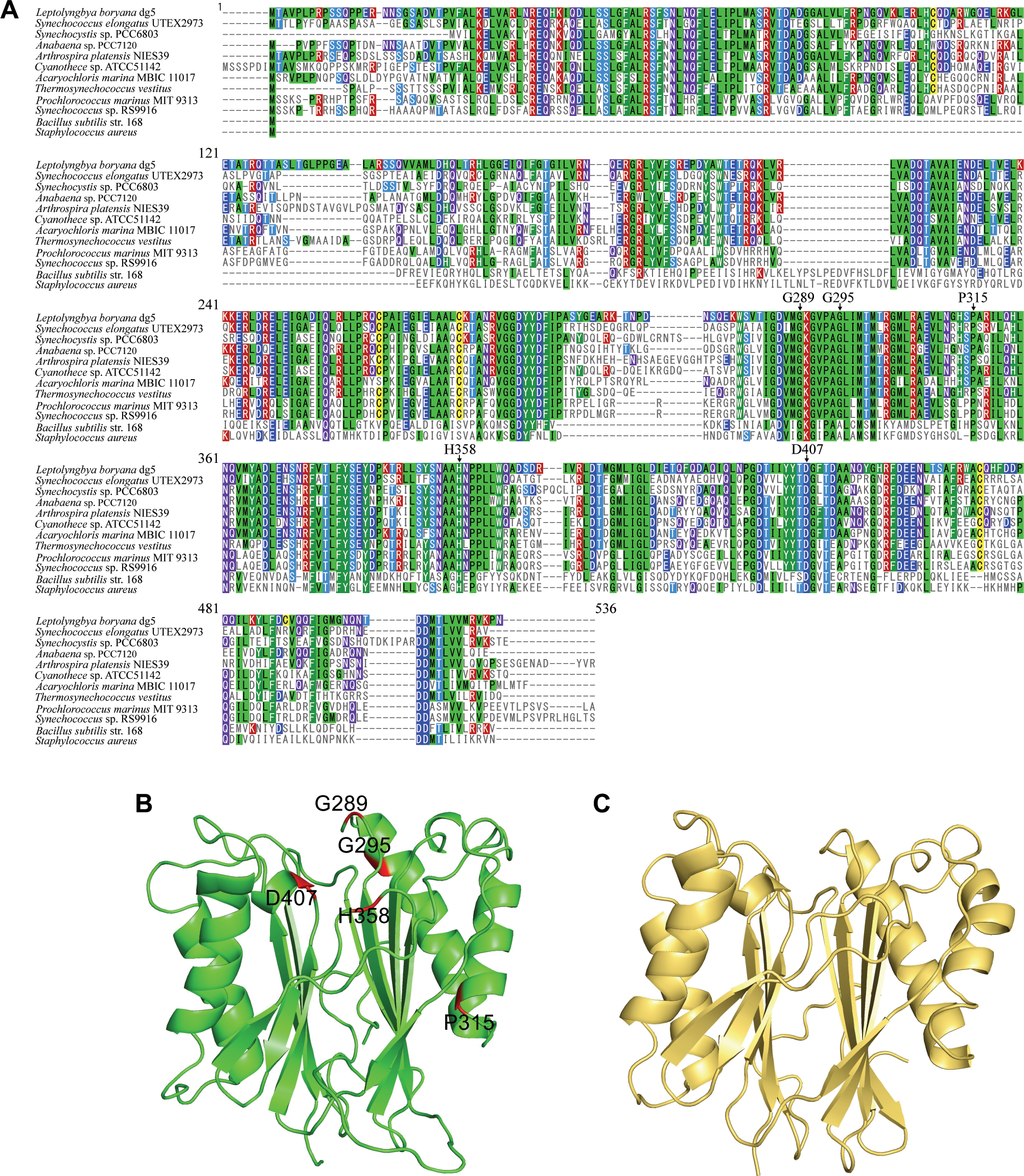
Multiple sequence alignment of RsbU/PhsP proteins from various organisms (**A**) and structural simulation of the C-terminal phosphatase domain of RsbU (**B**). (**A**) To examine the conservation of the amino acid sequence of the PhsP protein of *L. boryana*, multiple sequence alignment analysis was performed to compare the amino acid sequence information from other cyanobacterial and nonphotosynthetic bacterial species. Comparisons were made with *Synechococcus elongatus* UTEX 2973 (HltA, WP_039755543.1), *Synechocystis* sp. PCC 6803 (Slr2031), *Anabaena* sp. PCC 7120 (WP_010995926.1), *Arthrospira platensis* NIES 39 (WP_014274557.1), *Cyanothece* sp. ATCC 51142 (ACB49585.1), *Acaryochloris marina* MBIC11017 (WP_012164151.1), *Thermosynechococcus vestitus* (WP_011058070.1), *Prochlorococcus marinus* MIT 9313 (WP_011129391.1), *Synechococcus* sp. RS 9916 (WP_007099441.1), *Bacillus subtilis* str. 168 (WP_003234295.1), and *Staphylococcus aureus* (Q6XZ94). MAFFT (Katoh and Standley, 2016) was used for multiple sequence alignment. (**B, C**) 3D-structue prediction of PhsP of *L. boryana* using the MPI Bioinfomatics Toolkit (https://toolkit.tuebingen.mpg.de), HHpred (Soding et al., 2005), and MODELER. The structure of the RsbU proteins of *L. boryana* (**B**) was predicted from the known structure of the RsbU domain of MursiF (Rv1364c) of *M. tuberculosis* (PDB code:3KE6, **C**) using HHpred. The structure of the PhsP protein of *L. boryana*, which is highly homologous to HHpred, was predicted using MODELER. Residues with amino acid substitutions in *L. boryana* are shown in red.

Second, under dark-heterotrophic conditions, the expression of 932 genes was significantly altered by the presence or absence of LBDG_21500 (Fig. 6Bb, Fig. S7B). Most (28 genes) of the top 30 genes of the 932 genes that varied had significantly increased transcript levels (>Fig. 6Bb, Table S8). However, most genes were of unknown function. GSEA showed that the 547 genes that showed higher transcript levels in Δ21500 relative to dg5 were categorized in porphyrin metabolism, photosynthesis, biosynthesis of cofactors, galactose metabolism, and starch/sucrose metabolism (KEGG functional categories, Fig. S9B). However, it was difficult to infer how these increased transcript levels contributed to the promotion of dark-heterotrophic growth while sugar metabolisms could contribute.

A gene annotated as *psaK* (LBDG_37750) was found among the top 28 genes (Fig. 6Bb, Table S8). PsaK is a small subunit of the photosystem I complex and is located at the periphery of the PsaA side of the complex to bind two Chl molecules. In the photosystem I trimer structure, PsaK interacts with PsaB in the adjacent photosystem I unit (Toporik et al. 2020). The genome of *Synechocystis* sp. PCC 6803 encodes two *psaK* homologs, *psaK1* and *psaK2*, of which *psaK2* is involved in state transition under high-light conditions (Fujimori et al. 2005). In the genome of *L. boryana*, there are three *psaK* homologs, two of which appear to correspond to *psaK1* and *psaK2*. The third (LBDG_37750, *psaK3*) belongs to the divergent type PsaK (Fujimori et al. 2005), whose role is still unknown. It would be worthwhile to examine whether the increase in *psaK3* transcript levels in dark-grown Δ21500 contributes to the promotion of growth in the dark.

The three adjacent genes, *LBDG_46120*, *LBDG_46130*, and *LBDG_46140*, in the top 30 genes showed ratios (Δ21500-heterotroph/dg5-heterotroph) of 8.1, 10.7, and 6.7, respectively, indicating that their transcript levels are highly upregulated in Δ21500 grown under dark-heterotrophic conditions (Table S8). Although LBDG_46130 is a hypothetical protein lacking functional information, LBDG_46120 and LBDG_46140 are homologous proteins with a high similarity of 69.2%, which are both annotated as zinc-dependent alcohol dehydrogenases. This enzyme has recently been identified as a novel alcohol dehydrogenase through biochemical analysis of a probable ortholog in *B. subtilis* R5 (AMS48275 showing 45% and 43% homology to LBDG_46120 and LBDG_46140, respectively) (Ashraf et al. 2017), suggesting that the elevated transcript levels of these genes lead to increased alcohol dehydrogenase activity, which may contribute to the enhanced dark-heterotrophic growth in Δ21500 by increasing sugar fermentation activity. Another isoform, LBDG_35600, is also present in *L boryana* dg5, and its transcript level was increased 4.7-fold in Δ21500, suggesting that it is similarly upregulated. The contribution of the novel alcohol dehydrogenase to dark heterotrophic growth should be confirmed by isolating gene-knockout mutants in the future.

The RNA-seq data showed that the transcript levels of many genes were lower than those of dg5 under photosynthetic conditions due to the loss of *LBDG_21500*, suggesting that *LBDG_21500* contributes to the activation of the expression of many genes under photosynthetic conditions via the PSS involving LBDG_21500. Conversely, under dark-heterotrophic conditions, the transcript levels of many genes were increased in Δ21500; it is difficult to explain the increased transcript levels simply from the probable function of LBDG_21500 and the PSS. Loss of LBDG_21500 may lead to dysregulation of other transcriptional regulatory systems, such as other PSSs and two-component regulatory systems. For example, the transcript level of the response regulator (LBDG_46620) was greatly reduced in Δ21500 under photosynthetic conditions, and those of histidine kinases (LBDG_X2620, LBDG_X2630) and response regulators (LBDG_31480) were greatly increased in Δ21500 under dark-heterotrophic conditions.

Because loss of function of LBDG_21500 results in slower photosynthetic growth and faster heterotrophic growth than in the parental strains in *L. boryana*, with clearly different transcript profiles, we propose to rename LBDG_21500 as *phsP* (<u>p</u>hototrophic–<u>h</u>eterotrophic <u>s</u>witching <u>p</u>hosphatase).

## Discussion

In this study, using the high heterotrophic growth capacity of *L. boryana*, 28 dark-adapted variants were isolated through long-term cultivation under dark heterotrophic conditions, and 84 mutations were identified via genome resequencing analysis. Nineteen of them occurred in *phsP* (LBDG_21500), which encodes a PP2C-type serine phosphatase, and the phenotype of the targeted mutant Δ*phsP* (Δ21500) confirmed that *phsP* is the gene responsible for the common phenotype, decreased photosynthetic growth, and enhanced dark-heterotrophic growth, of the 19 variants.

### Conserved amino acid residues in PhsP

In *B. subtilis*, phosphatase RsbU, with which PhsP shows high similarity, has the activity to remove a phosphate group from the phosphorylated form (Ser56-P) of the anti-sigma antagonist RsbV in the PSS gene regulatory system in response to environmental stress (Yang et al. 1996). The dephosphorylated RsbV binds to the anti-sigma factor RsbW, which is bound to the sigma factor B, and the free sigma factor B recruits RNA polymerase to activate transcription, resulting in the induction of the transcription of stress response genes. The phosphatase activity of RsbU is activated by RsbT to integrate environmental signals into sigma B activation (Chen et al. 2003).

RsbU of *B. subtilis* is divided into an N-terminal and a C-terminal domain, and the N-terminal domain interacts with RsbT to activate the phosphatase activity of the C-terminal domain (Delumeau et al. 2004). PhsP of *L. boryana* (474 residues) can also be divided into an N-terminal (residues 1–221, containing a GAF domain) and a C-terminal (residues 222–474; Fig. 2C) phosphatase domain. Interestingly, all five amino acid substitutions in PhsP identified in dark-adapted variants occurred only in the C-terminal phosphatase domain. The amino acid sequence of PhsP was compared to putative orthologs in other cyanobacteria and to RsbUs in *B. subtilis* and *Staphylococcus aureus*. All five amino acid substitutions occurred at residues that are almost completely conserved in PhsP (G289, G295, P315, H358, and D407), except for G295, which is alanine only in *B. subtilis* and *S. aureus* (Fig. 8A). Functional analysis (Sachdeva et al. 2008) and the crystal structure of the RsbU domain of Rv1354c (MursiF) of *Mycobacterium tuberculosis* have been reported (Fig. 8C). Based on the structure of MursiF, we simulated the three-dimensional structure of PhsP and estimated the spatial positions of the five amino acid substitutions (Fig. 8B). The RsbU domain of MursiF exhibited Mn^2+^-dependent phosphatase activity. Four of the five amino acid residues, except Pro315, were presumed to localize around these metal ions. In particular, Asp407 (D407G) corresponded to one of the three conserved aspartic acids that coordinated the metal ions; the D328A substitution in MursiF resulted in a significant decrease in phosphatase activity (Sachdeva et al. 2008). This supports the hypothesis that substitutions of these residues detected in dark-adapted variants of *L. boryana* may result in a decrease or loss of phosphatase activity.

### A photosynthetic-heterotrophic switching mechanism

Given that the function of PP2C-type serine phosphatase in the PSS of other prokaryotes such as *B. subtilis*, PhsP functions as a phosphatase that dephosphorylates an anti-sigma antagonist in an unknown PSS regulatory system in *L. boryana*. Among the mutations other than *phsP* in the dark-adapted variants, mutations in *LBDG_16410*, encoding a putative anti-sigma antagonist, were identified in two variants (v2 and dg219). In dg219, a single-base insertion into the coding region caused a frameshift in *LBDG_16410*. In v2, a transposon insertion occurred in the intergenic region between *LBDG_16410* and of the adjacent *LBDG_16400* (encoding the carbamyl phosphate synthase small subunit), which may affect the transcript level of *LBDG_16410*. Both v2 and dg219 commonly exhibit the phenotype of loss of photosynthetic growth capacity. Because v2 and dg219 accompany mutations other than *LBDG_16410*, experimental confirmation is needed to determine whether the mutations in *LBDG_16410* are responsible for the loss of photosynthetic growth capacity. Furthermore, it is also important to examine whether the anti-sigma antagonist encoded by *LBDG_16410* is a dephosphorylation target of PhsP.

It remains unclear which sigma factor’s activity is regulated by the PSS involving PhsP. PhsP is presumed to function in this unknown PSS regulatory system via dephosphorylation of the anti-sigma antagonist to contribute to the optimization of the transcript profiles for various proteins involved in the balance between photoautotrophic and chemoheterotrophic metabolism in response to some environmental signals such as light and glucose.

As many as 20 genes encoding sigma factors exist in the *L. boryana* genome, which is significantly more than the 9 genes found in *Synechocystis* sp. PCC 6803 and the 12 found in *Anabaena* sp. PCC 7120 (Srivastava et al. 2020). This suggests that *L. boryana* has more complex sigma factor-mediated gene regulatory systems than these model cyanobacteria. Furthermore, two other genes (*LBDG_41040* and *LBDG_48270*) showed significant similarity (approximately 28%) to PhsP. An anti-sigma factor (LBDG_41050) is encoded immediately downstream of LBDG_41040. All amino acid residues (G289, P315, H358, and D407) which substituted in the dark-adapted variants, except for G295, are also conserved in the two PhsP-like proteins. Additionally, there are five genes for anti-sigma factors and seven genes for anti-sigma antagonists in the *L. boryana* genome. Further studies are needed to determine which anti-sigma antagonists, anti-sigma factors, and sigma factors operate with PhsP, and which environmental signals regulate global gene expression through PhsP.

In the model cyanobacterium *Synechocystis* sp. PCC 6803, a 154-bp DNA fragment corresponding to the N-terminal domain of *slr2031* (a *phsP* homolog) is deleted in a glucose-resistant strain (Katoh et al. 1995), and increased high-light tolerance and higher Chl and phycocyanin contents have been reported as phenotypes of the strain (Kamei et al. 1998). Additionally, a single *slr2031*-deficient mutant (Δ*slr2031*) was isolated from *Synechocystis* sp. PCC 6803, and the phenotype was reported (Huckauf et al. 2000). In the Δ*slr2031* mutant, Chl and phycocyanin contents were significantly increased compared with those in the WT. Conversely, in *L. boryana* Δ*phsP*, Chl and phycobilin contents were rather decreased under photosynthetic conditions, which appears inconsistent in these species. In the cyanobacterium *Synechococcus elongatus* UTEX 2973, which shows a very short doubling time for photosynthetic growth, an *rsbU* homolog plays an essential role in high-light tolerance, and the gene name *hltA* (high light tolerance A) was proposed (Walker and Pakrasi 2022). PhsP and HltA show high homology with each other (Fig. 8A). Interestingly, a single amino acid substitution, C35R, in the N-terminal domain of HtlA is the mutation responsible for the high-light sensitivity phenotype, which is in contrast to the case of PhsP in *L. boryana*, where all mutations were accumulated in the C-terminal domain.

The different phenotypes of *rsbU*-knockout mutants in different cyanobacterial species suggest that the PSS regulatory system, including the RsbU phosphatase, has evolved independently as a unique regulatory mechanism in each species. In particular, the high heterotrophic growth capacity that allows *L. boryana* to grow in the presence of glucose even in the dark may be supported by proper regulation (some suppressing regulation) via the PSS regulatory system involving PhsP, which could have evolved uniquely in the *L. boryana* lineage.

Protein–protein interaction network analysis using PhsP as the query detected anti-sigma antagonists (LBDG_08790, LBDG_52190) and an anti-sigma factor (LBDG_38100) (Fig. S10). Additionally, some sensor histidine kinases and response regulators of two-component regulatory systems were found to interact with candidate proteins. The two-component regulatory system, Hik33-RpaB, plays a critical role in proper photosynthetic growth in *Synechocystis* sp. PCC 6803 (Seino et al. 2009; Muramatsu and Hihara 2012; Riediger et al. 2019). During adaptation to various environments through switching between light-photosynthetic and dark-heterotrophic growth, many genes could be globally regulated by an elaborate and complex transcriptional regulatory network, in which PSSs, including PhsP, and two-component regulatory systems operate with extensive cross-talk in *L. boryana*.

Several dark-adapted variants obtained through long-term cultivation under dark heterotrophic conditions carried many mutations in genes involved in biosynthesis of photosynthetic pigments such as phycobilin and carotenoids, resulting in various colors such as khaki, cobalt blue, and yellow green. In addition to culture color alteration, other unique phenotypes such as abnormal cell morphology (dg204) and pigment leakage (v6) were observed. Such phenotypes are rarely selected under natural photoautotrophic conditions. *L. boryana* provides an excellent system for microevolutionary experiments leading to loss of photosynthesis.

## Experimental procedures

### Cyanobacterial strains and their cultivation

The filamentous cyanobacterium *Leptolyngbya boryana* IAM M-101 (WT) and dg5 (Fujita et al. 1996; Hiraide et al. 2015) were cultivated in BG-11 medium (Rippka et al. 1979) supplemented with 20 mM HEPES-KOH (pH7.4) under continuous light (20–25 μmol_photon_ m^−2^ s^−1^; FL15D fluorescent lamp, Panasonic, Osaka, Japan) under photoautotrophic conditions. For mixotrophic and heterotrophic conditions, cyanobacterial cells were grown in BG-11 medium containing 30 mM glucose (BG-11Glc) in the light and in the dark, respectively. Kanamycin (15 μg ml^−1^) and chloramphenicol (10 μg ml^−1^) were added to the cultures of mutants to maintain the drug marker gene.

### Isolation and maintenance of dark-adapted variants

To isolate the dark-adapted variants, the parent strains dg5 and WT were inoculated onto a BG-11 agar plate of containing 30 mM glucose and grown in the dark for 9 days and 40 days, respectively. Dense colonies that appeared on the plates were isolated as the first generation of dark-adapted variants (Fig. S1). The dark-adapted variants were then inoculated into a new BG-11 agar plate containing 30 mM glucose every 10 days to 2 weeks.

### Construction of plasmids

A plasmid for isolating Δ21500 was prepared as follows: DNA fragments of approximately 1.5 kb upstream and downstream of LBDG_21500 required for homologous recombination were amplified via polymerase chain reaction (PCR; PrimeSTAR Max DNA polymerase, Takara, Shiga, Japan) using *L. boryana* dg5 genomic DNA as a template. The kanamycin resistance cartridge (Tsujimoto et al. 2014) and vector (pYCSFX+SacB, pYCSFX (Fujita and Bauer, 2000) carrying the *sacB* gene of pZJD29a (Masuda and Bauer, 2004)) were also amplified separately via PCR. These four fragments (vector, upstream fragment, kanamycin resistance cartridge, and upstream fragment) were connected using the In-Fusion reaction (In-Fusion HD Cloning Kit, Takara, Shiga, Japan) to construct the LBDG_21500-disrupting plasmid (Table S9).

To construct the plasmid p21500_complement for complementation of Δ*21500*, the genomic region from approximately 1.5 kb upstream of *LBDG_21500* to 200 bp downstream of the termination codon was amplified via PCR using dg5 genomic DNA as the template. The other DNA fragment on the vector side was amplified via PCR using pZJD29a as a template, and these two fragments were connected using the In-Fusion reaction to construct a complementation plasmid.

Two variants of plasmids to examine site-directed variants, G295D and D407G, were constructed via site-directed mutagenesis using the PrimeSTAR Mutagenesis Basal Kit (Takara) with p21500_complement as the template. In G295D, the G at 884 bp from the start codon of *LBDG_21500* was replaced with A. In D407G, A at 1,220 bp from the start codon of *LBDG_21500* gene was replaced with G (Fig. S4). Each base substitution was confirmed using the Sanger method after plasmid construction.

### Transformation of L. boryana through conjugation

Transformation of *L. boryana* was performed using a conjugation method. The donor *Escherichia coli* S17-1 λ*pir* carrying the LBDG_21500-disrupting plasmid was prepared as described previously (Tomatsu et al. 2018). *L. boryana* dg5 cells grown photosynthetically on a BG-11 agar plate were suspended in 0.3 mL of BG-11 medium at 0.5 OD_730_ and mixed with the same volume of the donor *E. coli*. A sterile nylon membrane (Hybond-N+; 82 mm, GE Healthcare, Chicago, IL) was placed on a BG-11 agar plate without antibiotics, and the cell suspension was spread on the membrane and allowed to dry for 5 min. The agar plate was incubated under photosynthetic conditions (20–25 μmol_photon_ m^−2^ s^−1^) at 30°C for 24 h. The membrane was transferred to a BG-11 agar plate with appropriate antibiotics and incubated further under photosynthetic conditions. Drug-resistant colonies that appeared after approximately 7–10 days were isolated as transconjugants. The isolation of transconjugants for complementation analysis was completed at this stage.

In the case of the isolation of Δ21500, to remove the WT LBDG_21500 copy and the vector carrying *sacB* by the second homologous recombination, the first isolated transconjugants were spread on BG-11 agar medium supplemented with 5% sucrose and appropriate antibiotics, and incubated for 7–10 days to allow the appearance of sucrose-resistant colonies. However, the LBDG_21500-null mutant was difficult to isolate under normal photosynthetic conditions. Thus, the first recombinants were grown under mixotrophic conditions under low-light conditions (5 μmol_photon_ m^−2^ s^−1^) in the presence of 30 mM glucose, resulting in the successful isolation of the mutant, Δ*21500*. Knockout of the target gene was confirmed via colony PCR (Quick Taq® HS DyeMix, Toyobo, Osaka, Japan).

### Measurement of the absorption spectra of cell suspensions

Cell suspensions were adjusted to 0.1 OD_730_ (model UV-1600, Shimadzu, Kyoto, Japan), and their absorption spectra were measured (model V-650; Jasco, Hachioji, Japan) using an integrating sphere unit (ISV-722; Jasco).

### Determination of Chl content

Cells were suspended in 90% (v/v) methanol, placed on ice for 1 h, centrifuged at 15,000 rpm for 20 min at 4°C, and the absorption spectrum of the supernatant was measured using a spectrophotometer (model V-650). The Chl concentration was calculated using the formula (Tandeau de Marsac and Houmard 1988) or molar absorption coefficient for Chl (Porra et al. 1989).

### Carotenoid pigment analysis

Cells were lysed in acetone/methanol (7:2, v/v) using an ultrasonicator for 20 s at 4°C, and the organic fraction was obtained via centrifugation at 15,000 rpm at 4°C for 10 s. After evaporation to dryness, the pigments were dissolved in a small volume of chloroform/methanol (3:1, v/v). An HPLC system equipped with a 2998 pump (Waters, Milford, MA) and a SPD-M10A photodiode array detector (Shimadzu, Kyoto, Japan) was used. For the analysis of total extracted pigments, a µBondapak C_18_ column (100 × 8 mm, RCM-type: Waters) was used. A linear gradient from methanol/water (9:1, v/v) to methanol was applied for 20 min, followed by methanol elution at a flow rate of 1.8 mL min^−1^ (Fig. 3A-D; Takaichi et al. 2020). For the analysis of carotenes, total pigments were dissolved in a small volume of hexane, and then loaded on a silica gel 60 column (Merck, Darmstadt, Germany). Carotenes were eluted with hexane, whereas xanthophylls and Chl remained on the column. A NOVA C_18_ column (100 × 8 mm, RCM-type; Waters) was used, and acetonitrile/methanol/tetrahydrofuran (58:35:7, v/v) was used for isocratic elution at a flow rate of 1.8 mL min^−1^ (Fig. 3F, G; Takaichi, 2000). To identify the pigments, the retention times on HPLC and absorption spectra of the pigments were compared with those from *Anabaena* sp. PCC 7120 (Takaichi et al. 2005).

### Preparation of the thylakoid membrane

Cells grown on an agar plate under photoautotrophic conditions for 3 days were incubated in 3 mL of thylakoid buffer (50 mM HEPES-NaOH; pH 7.0, 10 mM MaCl_2_, 5 mM CaCl_2_, 25% (w/v) glycerol), 2 mM PMSF, and 100-mg glass beads (150–212 microns, Sigma-Aldrich, St. Louis, MO). The cell suspension was vortexed for 30 s and immediately placed on ice for 30 s; this was repeated five times, then the cell suspension was centrifuged at 3,000 rpm at 4°C for 5 min. Next, the supernatant was centrifuged at 40,000 rpm for 30 min at 4°C. The pellet was suspended in 500 μL of thylakoid buffer and stored at –80°C as the thylakoid membrane fraction. The protein concentration in the thylakoid membrane fraction was determined via Bradford’s method (Protein Assay, Bio-Rad, Hercules, CA) using a bovine serum albumin solution of known concentration as a standard.

### SDS-PAGE and Western blot analysis

The thylakoid membrane fraction prepared above was mixed with an equal volume of sample buffer (40 mM dithiothreitol, 5.2 % lithium dodecyl sulfate, 172 mM Tris, 0.5 M sucrose, and 0.01 % bromophenol blue), and samples were heat treated at 70°C for 10 min. Samples without heat treatment were also prepared. For Western blot analysis, samples with 1 μg of protein or 0.05 μg of Chl were loaded into each lane for SDS-PAGE. Electrophoresis was performed for 90 min at a constant voltage of 200 V using 6 M urea, 12 % homogeneous acrylamide separating gel and concentrated gel. Western blot analysis was carried out essentially as described (Yamazaki, 2006) and reacted with antisera against D1 (Agrisera, Vännäs, Sweden), D2 (Agrisera), CP47 (Kada et al. 2003), 33 kDa (Kada et al. 2003), PsaA (Agrisera), and cytochrome *f* (Agrisera). The specific protein bands were visualized with a secondary antibody (anti-rabbit IgG-HRP conjugate; Bio-Rad) using a chemiluminescent substrate (Amersham ECL Western Blotting Detection Reagents; GE Healthcare, Chicago, IL) with a lumino-image analyzer (iBrightTM FL1500 Imaging System; Thermo Fisher Scientific, Waltham, MA).

### Preparation of genomic DNA

Cells were grown on an agar plate (9 cm in diameter) under photosynthetic conditions for approximately 14 days, and cells on half the area of the agar plate were collected and suspended in 500 μL of TE buffer (pH 8.0) and freeze-thawed three times using liquid nitrogen. Cell lysis and subsequent preparation of genomic DNA were performed as described in Uesaka et al (2024).

### Preparation of a library for genome resequencing

Concentrations of the extracted genomic DNA samples were accurately measured using the fluorescence measurement method by binding of SYBR Green (Ex 485 nm, Em 535 nm). DNA solutions of approximately ≥5 ng μL^−1^ were subjected to the tagmentation reaction using the Nextera DNA Flex Library Prep Kit (Illumina, San Diego, CA) was used at 1/40 scale of the provided protocol. An aliquot (15 ng; 1.5 μL) of DNA solution was mixed with 1.0 μL of reaction master mix, and the reaction was performed in a thermal cycler at 55°C for 25 min and 80°C for 2 min. Thereafter, 100 μL of Tagment Wash Buffer was added, and the beads in the reaction solution were trapped with a magnet to wash off the supernatant containing unreacted genomic DNA and transposase that had been removed via thermal denaturation.

The PCR master mix (20 μL) that was prepared using KAPA HiFi HotStart Ready Mix (Roche, Basel, Switsland) (×2 KAPA HiFi PCRmix, 12.5 μL; milliQ water, 7.5 μL) was added to each well of the PCR plates and mixed well (2,000 rpm, 1 min). A barcode was assigned to each well of the PCR plate, and 2.5 μL of two types of adapters (i7XX and i5XX) were added to each well. PCR was performed in a thermal cycler under the following conditions: initial reaction at 72°C for 3 min and 98°C for 30 s, followed by 12 cycles of denaturation (98°C for 30 s), annealing (62°C for 15 s), and extension (72°C for 2 min), followed by a final reaction at 72°C for 1 min.

The PCR-reacted samples were trapped using a magnet, and 45 μL of the supernatant was transferred to another PCR plate. AMPure XP bead suspension (13.5 μL) was added, and the plate was agitated at room temperature for approximately 5 min. Subsequently, it was left on the magnetic rack for at least 2 min, and then 58.5 μL of the supernatant was transferred to a new plate after confirming that the magnetic beads were trapped in the rack. To this, 15.8 μL of AMPure XP bead suspension was added, and the suspension was vortexed at room temperature for approximately 5 min. The suspension was placed on a magnetic rack again for at least 2 min, and the supernatant was discarded after confirming that the magnetic beads were trapped in the rack. With the beads trapped, 200 μL of 80% ethanol solution was added, the suspensions were allowed to stand for a while, and then the ethanol solution was removed. This was repeated once more. The plate was removed from the magnetic rack and centrifuged to remove the ethanol solution completely. Water (30 µL) was added to the plate, mixed well, and the tube was allowed to stand for at least 1 min, and then the supernatant (25 µl) was transferred it to a new plate.

DNA concentrations were measured and recorded using fluorescence (SYBR Green). More than 10 ng of DNA was subjected to MultiNA (Shimadzu) to measure the DNA size distribution. The actual libraries were selected to be within a certain molecular weight distribution (300–1000 bp) using AMPure XP beads. The concentration of the libraries was corrected from the weight concentration determined through fluorescence measurement using the molecular weight estimate from the MultiNA analysis. All samples were mixed and purified again with AMPure XP beads. Finally, library samples were sent to Macrogen Japan (Tokyo, Japan) for genome resequencing (HiSeqX). The number of reads obtained via sequencing is shown in Tables S4 and S5.

### Variant calling for dark-adapted variants

For quality and adapter trimming of sequencing reads, the fastp preprocessor (Chen et al. 2018) was used with the condition “-q 15 -n 10 -t 1 -T 1 -l 20.” The remaining sequencing reads were used for further analysis. The statistics for raw and trimmed sequencing reads are shown in Tables S5 and S6. Mutation calls were performed using breseq (Deatherage and Barrick 2014) with the consensus mode (settings for isolated/cloned samples) for base substitution (hereafter SNV), small insertion and deletion (hereafter, small indel), and large deletion. The reference genome was *L. boryana* IAM M101 (AP014638, AP014639, AP014640, and AP014641; Hiraide et al. 2015). All positions called using breseq were visually checked using the IGV (Integrative Genomics Viewer; Thorvaldsdóttir et al. 2013) genome browser. Those with a mutation rate of 100% (+) were listed, and if a nonpure mutation was detected at the same locus in another strain, it was denoted with a “+/-.” Then, using the breseq utility tool gdtools, we created a table that integrated the mutations for each sample.

Although breseq does not support the detection of structural mutations (e.g., transposon insertions), we considered an abnormal mapping pattern of paired-end reads as an indication of a mutation and added an “Unassigned new junction evidences” entry to the table (see the breseq online manual; https://barricklab.org/twiki/pub/Lab/ToolsBacterialGenomeResequencing/documentation/).

The loci called in this field were checked on IGV to determine if there were consecutive mismatches in the read/genome alignment. If there is a series of mismatches in the alignment, it is expected that the read/genome alignment is completely incorrect. If such a read was identified, the sequence of the read was extracted and BLAST-searched against the reference genome. If an exact match was identified from a different locus in the genome, it was assumed to be the original site of the insertion sequence and added to the mutation table in breseq.

### Activities of photosynthesis and respiration

Cells were cultivated in liquid medium aerated with 2% (v/v) CO_2_ for 3 days under either photoautotrophic, photomixotrophic, or dark-heterotrophic conditions. The cells were collected via centrifugation and suspended in BG-11 liquid medium under the photoautotrophic conditions or in BG-11+Glc liquid medium under photomixotrophic and dark heterotrophic conditions. Oxygen evolution and consumption were measured in the cell suspension (1,980 µL) after adding 20 µL of NaHCO_3_ (final concentration, 10 mM) with an oxygen electrode (Oxytherm+, Hansatech, Norfolk, UK). After 4 min of illumination (50 μmol_photon_ m^−2^ s^−1^), the rate of oxygen evolution was measured. The electrode cuvette containing the cell suspension was incubated in the dark covered by a piece of black cloth for 4 min to measure the rate of oxygen consumption. After the measurements, the exact Chl concentration in the cell suspension was determined. The oxygen evolution and consumption rates per 1 μg _Chl_ mL^−1^-suspension and per min were calculated from a 2-min record, during which the rate of increase or decrease in oxygen concentration was constant in each measurement, and these measurements were used as the photosynthesis and respiration activities, respectively.

### Low temperature fluorescence spectra

Cells were cultivated under photoautotrophic conditions for 2 days in liquid medium aerated with 2 % (v/v) CO_2_. The cells were suspended in BG-11 medium at 5 μg _Chl_ mL^−1^, and the low-temperature fluorescence spectra were measured using a fluorometer (FP777w; Jasco) equipped with a cooling unit (PMA-281; Jasco).

### RNA preparation

Cells were grown on agar medium for 3 days under photomixotrophic conditions and then collected and suspended on BG-11 liquid medium for photoautotrophic and on BG-11Glc liquid medium for dark-heterotrophic cultivations. After cultivation for 2 days under the respective conditions, 50 mL of cell suspension was centrifuged (R12A2; CR21N, Eppendorf Himac Technologies, Hitachinaka, Japan) at 10,000 rpm at 4°C for 10 min, and the supernatant was removed. The cell pellet was suspended in 500 μL of TE buffer (pH 8.0). The cell pellets were suspended in 500 μL of TE buffer (pH 8.0), frozen and thawed twice with liquid nitrogen, incubated with 100 μL of 50 mg mL^−1^ lysozyme for 1 h at 37°C, and centrifuged at 8,000 rpm at 25°C for 1 min (AR015-24; MX-301; Tommy Seiko, Tokyo, Japan), following which the supernatant was removed. The resulting cell pellets were used to extract RNA using TRIzol® reagent (Thermo Fisher Scientific) and NucleoSpin RNA (Macherey-Nagel, Düren, Germany). Extracted RNA was quantified using a Nanodrop ND 1000 spectrophotometer (Thermo Fisher Scientific).

The RNA prepared as described above was sent to Macrogen Japan for library preparation and cDNA sequencing as follows: ribosomal RNA (rRNA) was removed from each sample using the NEBNext Bacteria rRNA Depletion Kit (New England BioLabs, Ipswich, MA), and library preparation was performed using the TruSeq Stranded Total RNA Library Preparation Kit (Illumina). Libraries were paired-end sequenced at 150-bp using the NovaSeq6000 (Illumina).

### RNA-seq data analysis

The statistics for raw sequencing reads are shown in Table S6. Raw sequencing reads were adapter trimmed and quality filtered using fastp. Preprocessed sequencing reads were mapped to a reference genome sequence (accession ID: GCA_002142495.1; Hiraide et al. 2015) using Bowtie2 v2.4.5 with the sensitive setting, and subsequently converted from the sam to bam format using SAMtools v.1.15. The number of sequencing reads aligned to each open reading frame was counted using featureCounts v.2.0.1 with the option “-s 0 -t exon -g gene_id.” To examine the data characteristics, the raw expression count was transformed to log2 CPM (count per million) after adding pseudocounts of “4.” The resulting matrices were analyzed using PCA and a hierarchical clustered heatmap to assess the quality between samples. The heatmaps and PCA plots were rendered using iDEP (integrated Differential Expression and Pathway analysis v0.95; http://bioinformatics.sdstate.edu/idep/). Raw count normalization and subsequent differential expression gene analysis were performed using Bioconducter’s DESeq2 package v.1.2.0 (Love et al. 2014) at a significance level of FDR < 0.01. GESA against KEGG functional categories was performed using Bioconductor’s clusterProfiler package v.4.4.3 (Wu et al. 2021). Volcano plots were rendered using the ggVolcanoR shiny server (https://ggvolcanor.erc.monash.edu) (Mullan et al. 2021). Each command was executed under default conditions unless otherwise specified.

## Acknowledgments

We thank Shinji Masuda for providing pJZD29a. We thank Tatsuhiro Tsurumaki for providing valuable suggestions on the analysis of photosystems, Yukako Hihara for providing suggestions on the categorization of RsbU, Asako Segawa and Mayu Chikada for offering technical assistance, Kazuki Terauchi for engaging in discussion; and Takafumi Yamashino, Hisanori Yamakawa, Natsuki Tanaka and all members of the Laboratory of Molecular and Functional Genomics for engaging in discussions and offering technical assistance.

## Author Contributions

Yuichi Fujita: Conceptualization; data curation; format analysis; investigation; supervision; resources; project administration; supervision; visualization; writing - review and editing; writing - original draft preparation; funding acquisition. Shintaro Hida: Data curation (equal); format analysis; investigation (equal); visualization; writing - review and editing. Marie Nishio: Data curation (equal); format analysis; investigation (equal); visualization; writing - review and editing. Kazuma Uesaka: Data curation (equal); format analysis; investigation (equal); software; visualization; writing - review and editing. Mari Banba: Investigation; resources; writing - review and editing. Nobuyuki Takatani: Investigation; writing - review and editing. Shin-ichi Takaichi: Investigation; writing - review and editing. Haruki Yamamoto: Investigation; supervision; writing - review and editing. Kunio Ihara: Investigation; Software; writing - review and editing.

## Data Availability Statement

Raw sequencing reads have been deposited in the DDBJ Sequence Read Archive (DRA) under BioProject accession number PRJDB15833; BioSample accession numbers SAMD00611845–SAMD00611873; and DRA accession numbers DRR469493–DRR469521. Raw RNA-seq sequencing reads have been deposited in the DDBJ DRA under BioSample accession numbers SAMD00611726–SAMD00611737 and DRA accession numbers DRR469481–DRR469492. Data are contained within the article or Supplementary Material. All raw data have been deposited into the data management system of Nagoya University.

