## Supplementary Figures S1 to S10 for "Microevolution toward loss of photosynthesis: Mutations promoting dark-heterotrophic growth and suppressing photosynthetic growth in cyanobacteria"

**Figure S1.**	Isolation of dark-adapted variants from dg5 and WT.

**Figure S2**.	Initial characterization of dark-adapted variants v1–v6.

**Figure S3.**	Initial characterization of dark-adapted variants dg201-dg229.

**Figure S4.**	Construction of a *LBDG_21500* knock-out mutant ∆21500, mutants carrying a point mutation in the LBDG_21500, and complementation with the WT copy of *LBDG_21500*.

**Figure S5.**	SDS-PAGE and Western blot analysis of *∆21500* and dg5 grown under photoautotrophic conditions.

**Figure S6.**	Principal component analysis of the CPM normalized read count of 12 samples.

**Figure S7.**	Volcano plots of the significantly differential expression genes by the knock-out of *LBDG_21500* under light and dark conditions, and in response to light and dark conditions of dg5 and *∆21500*.

**Figure S8.**	Venn diagrams showing number of genes with differential expressed genes (DEGs) under each condition.

**Figure S9.**	The results of gene set enrichment analysis (GSEA).

**Figure S10.**	Protein-protein interaction (PPI) network analysis of PhsP related proteins.

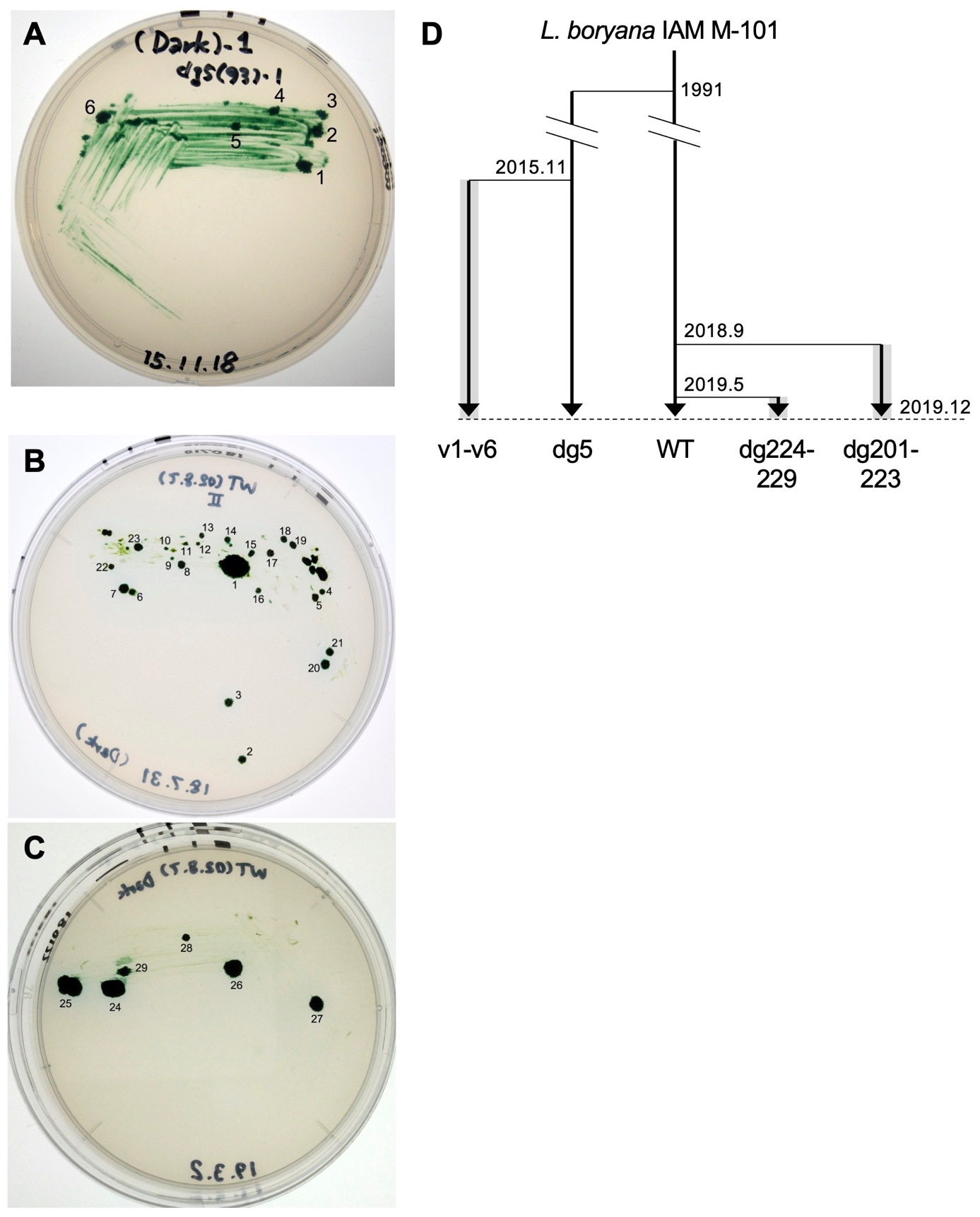

**Figure S1.**	Isolation of dark-adapted variants from dg5 (**A**) and WT (**B, C**). (**A**) Numbers 1–6 correspond to v1–v6, respectively. **B, C.** Numbers 1–29 correspond to dg201–dg229, respectively. (**D**) Brief history of the variants. Dark-heterotrophic cultivation is shown shaded in gray. Seven colonies corresponding to the missing numbers were discarded without continued cultivation because they did not grow well under dark-heterotrophic conditions after pickup.

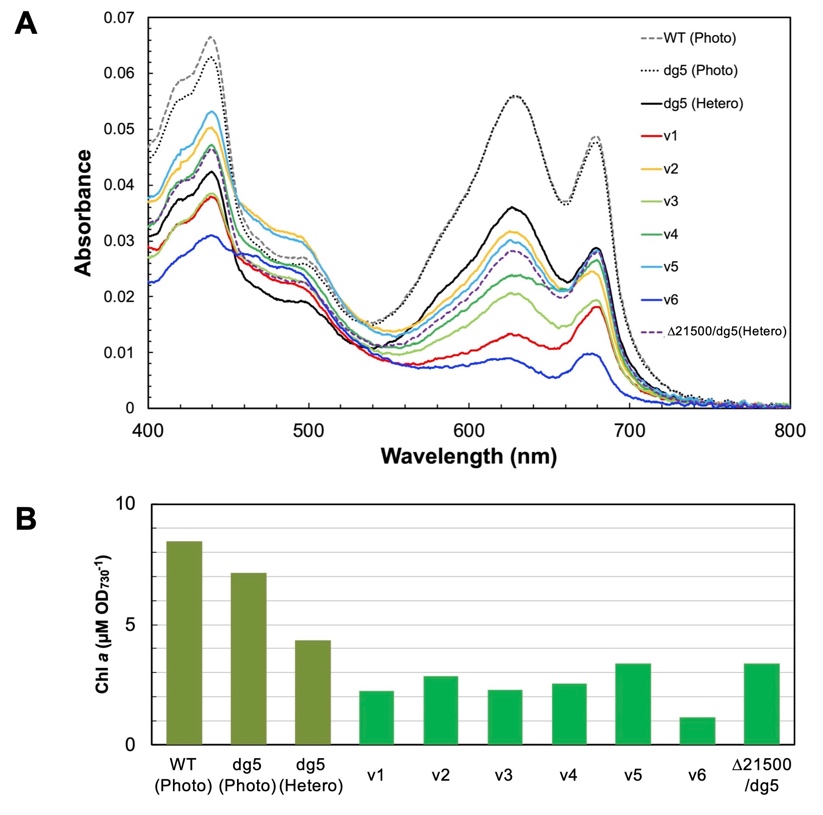

**Figure S2.**	Initial characterization of dark-adapted variants v1–v6. (**A**) Absorption spectra of dark-grown cells of v1–v6. As controls, absorption spectra of WT and dg5 cells grown under photoautotrophic conditions are also shown. (**B**) Chlorophyll contents of dark-grown cells of v1–v6.

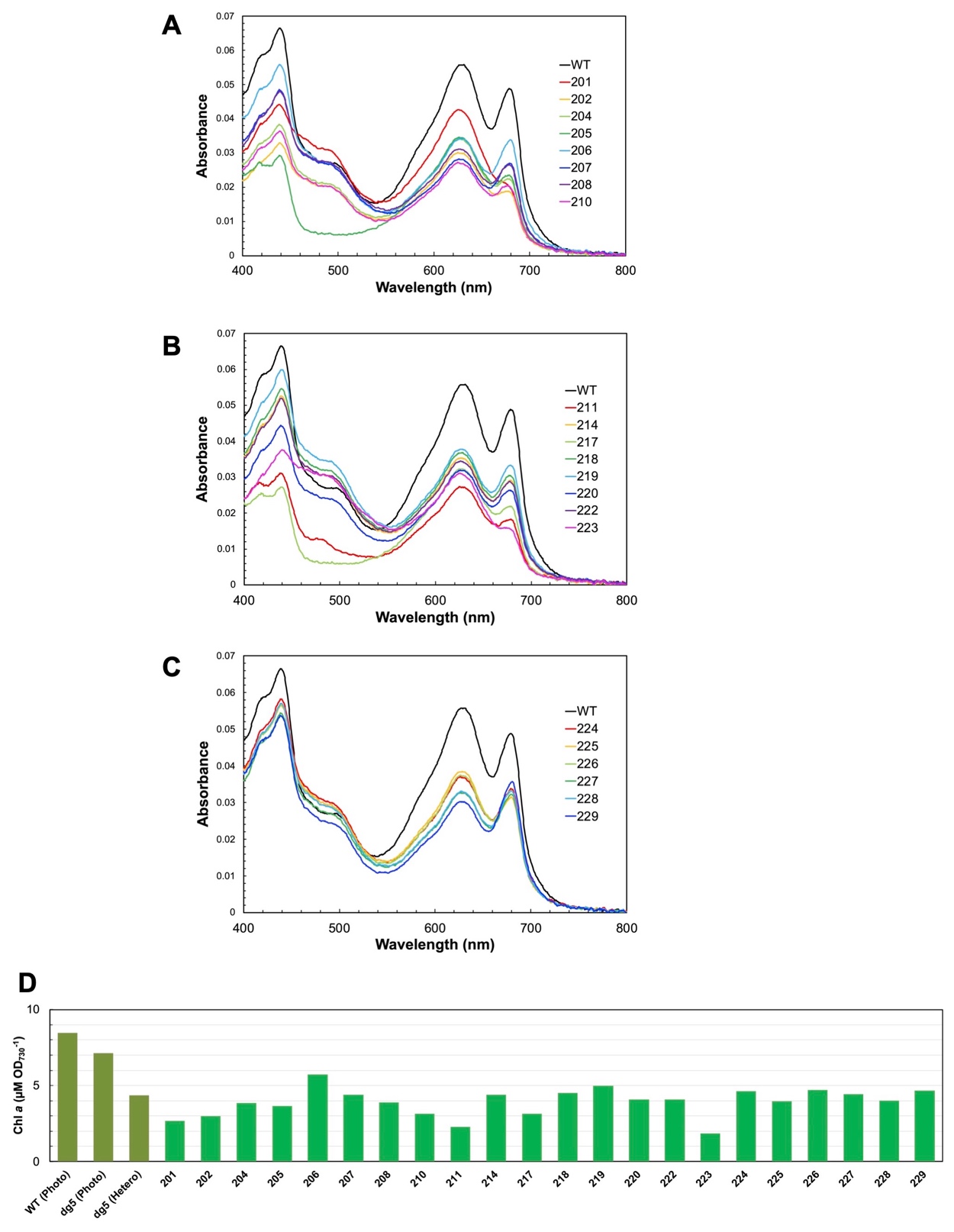

**Figure S3.**	Initial characterization of the dark-adapted variants dg201–dg229. (**A-C**) Absorption spectra of dark-grown cells of dg201–210 (**A**), dg211–dg223 (**B**), and dg224–229 (**C**). As controls absorption spectra of WT and dg5 cells grown under photoautotrophic conditions are also shown. (**D**) Chlorophyll contents of dark-grown cells of dg201–dg229.

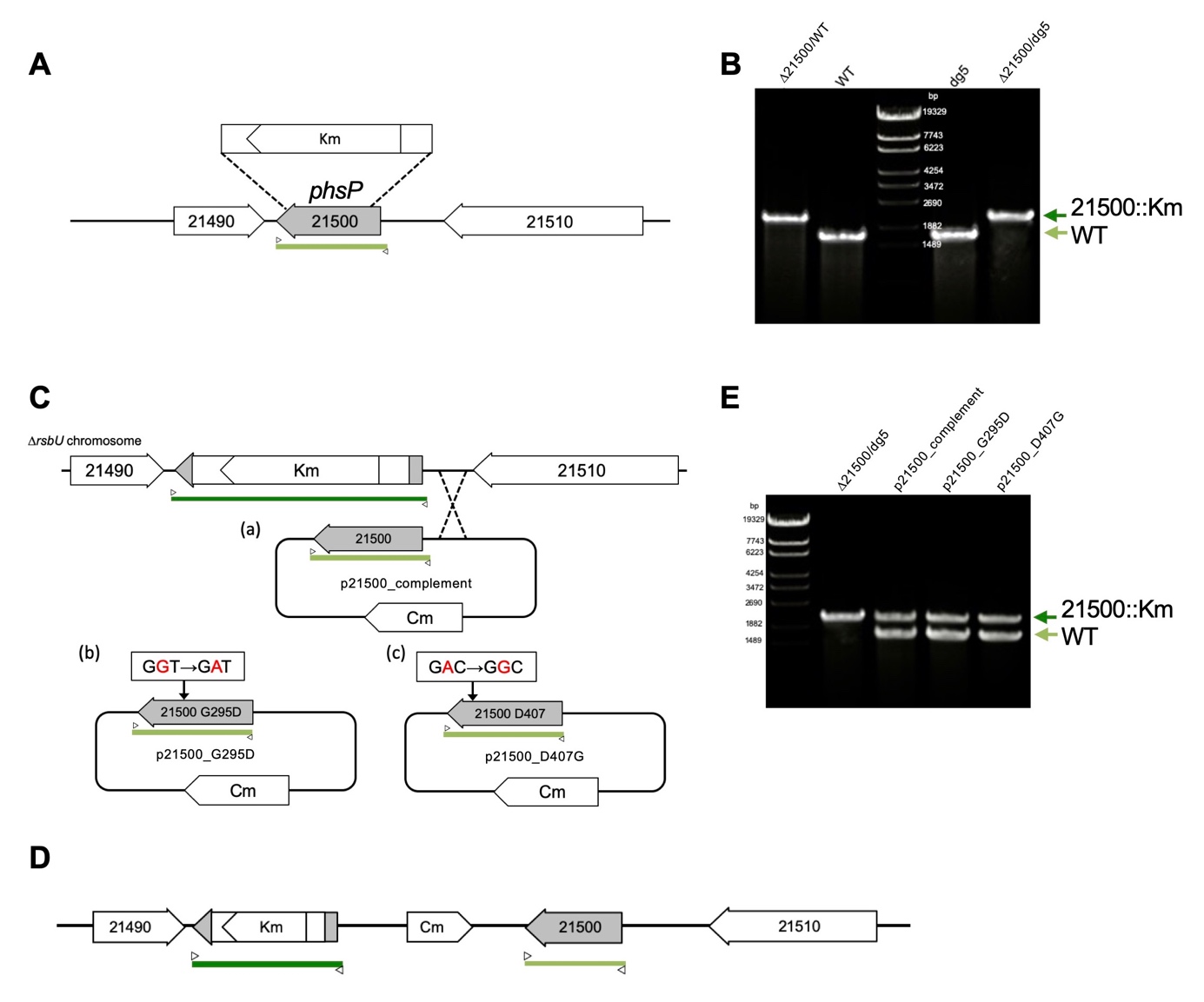

**Figure S4.**	Construction of a *LBDG_21500* knock-out mutant, ∆21500 (**A**, **B**), mutants carrying a point mutation in the LBDG_21500 (**C**, **E**), and complementation with the WT copy of *LBDG_21500* (**D**, **E**). (**A**) Isolation of *∆rsbU* mutants. Most of the coding region of *LBDG_21500* was replaced with the kanamycin resistance gene (Km). The chromosomal region amplified by PCR is indicated by the yellow green bar. (**B**) Confirmation of the complete loss of the WT copy of the *LBDG_21500* gene by PCR. (**C**) Complementation with the WT copy (a) or *LBDG_21500* variants (G295D (b); D407G (c)). The plasmid carries the chloramphenicol resistance gene (Cm). (**D**) Chromosomal region of *LBDG_21500* in the complementation mutants. The chromosomal regions for the WT copy and knockout copies amplified by PCR are shown by the yellow, green, and green bars, respectively. (**E**) Confirmation of the complementation of *LBDG_21500* in the transformants. Primers used in PCR are indicated by small white triangles.

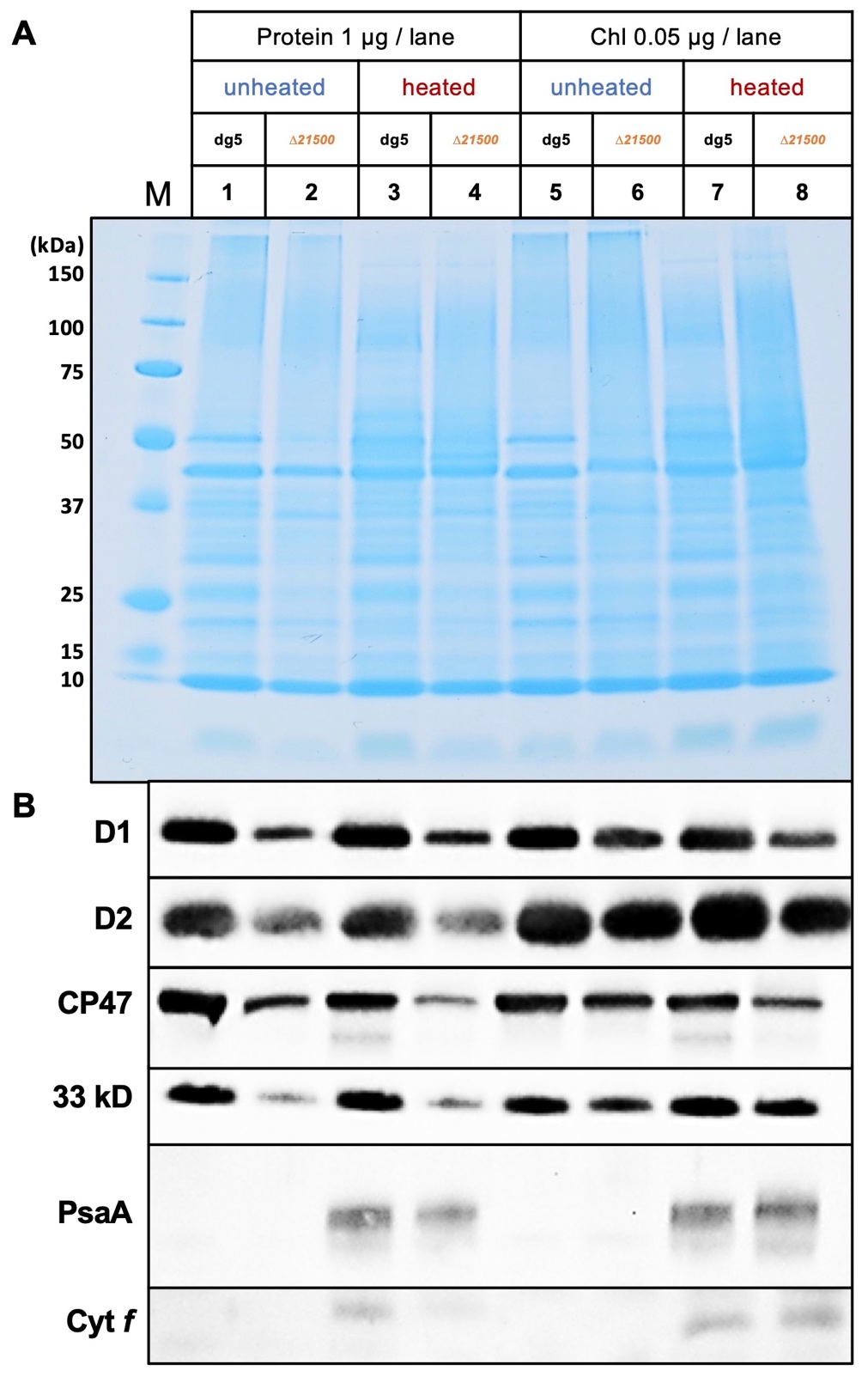

**Figure S5.**	SDS-PAGE (**A**) and western blot analysis (**B**) of *∆21500* (lanes 1, 3, 5 and 7) and dg5 (lanes 2, 4, 6 and 8) grown under photoautotrophic conditions. Samples were loaded onto each lane with a protein (lanes 1–4) or Chl (lanes 5–8) base. Protein samples were unheated (lanes 1, 2, 5, and 6) or heated (lanes 3, 4, 7 and 8). After electrophoresis, the protein signals were stained with Coomassie brilliant blue (Quick-CBB Plus, Wako) (**A**). Specific protein signals were detected using specific antisera against D1, D2, CP47, 33 kD (PsbO), PsaA and cytochrome *f* (**B**). Panel B is identical to Figure 5C.

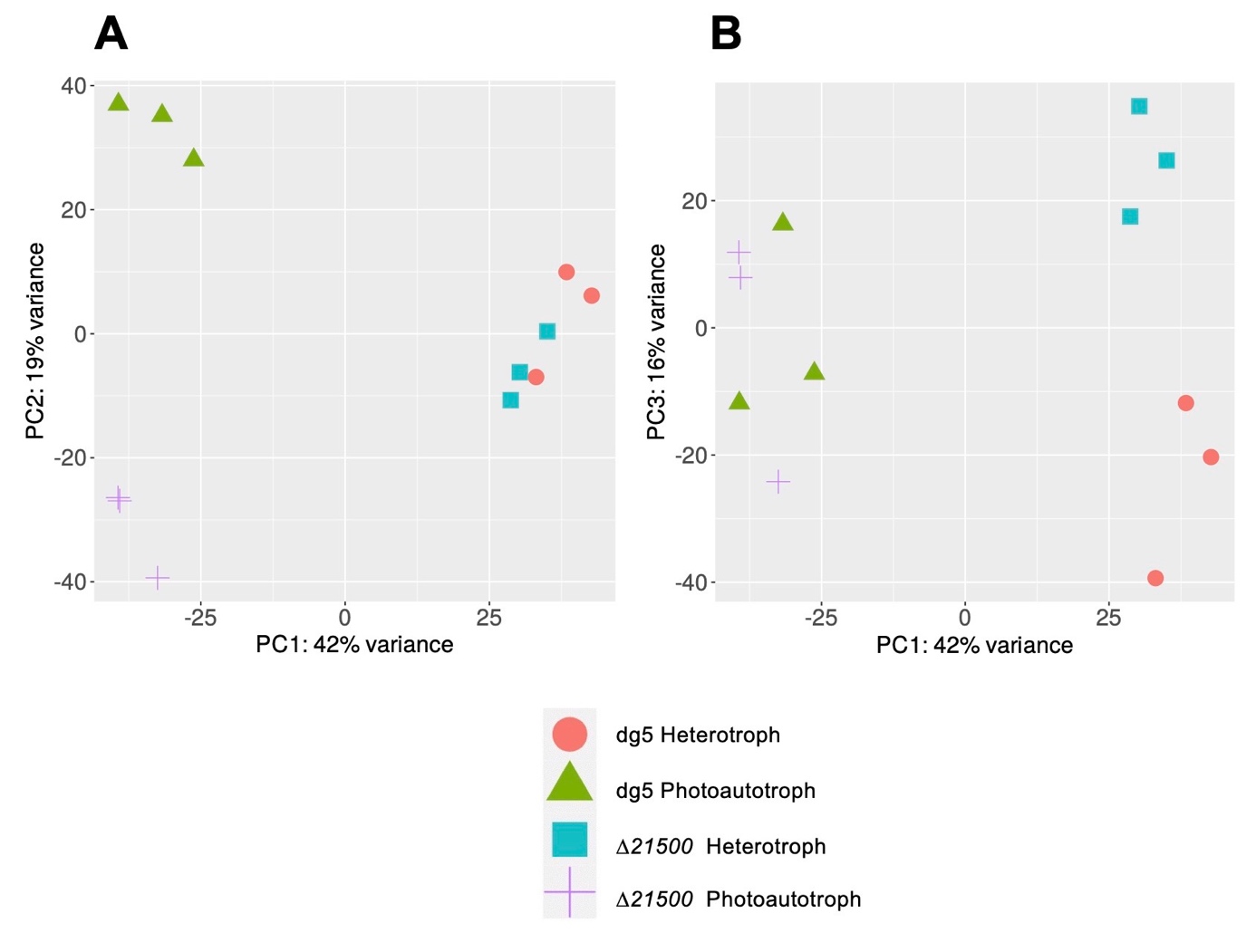

**Figure S6.**	Principal component analysis of the count per million (CPM)-normalized read count of 12 samples. Principal component analysis of the first and second principal components (**A**), which accounted for 61% of the variation, and the first and third principal components (**B**), which accounted for 58% of the variation. Plots are colored according to conditions.

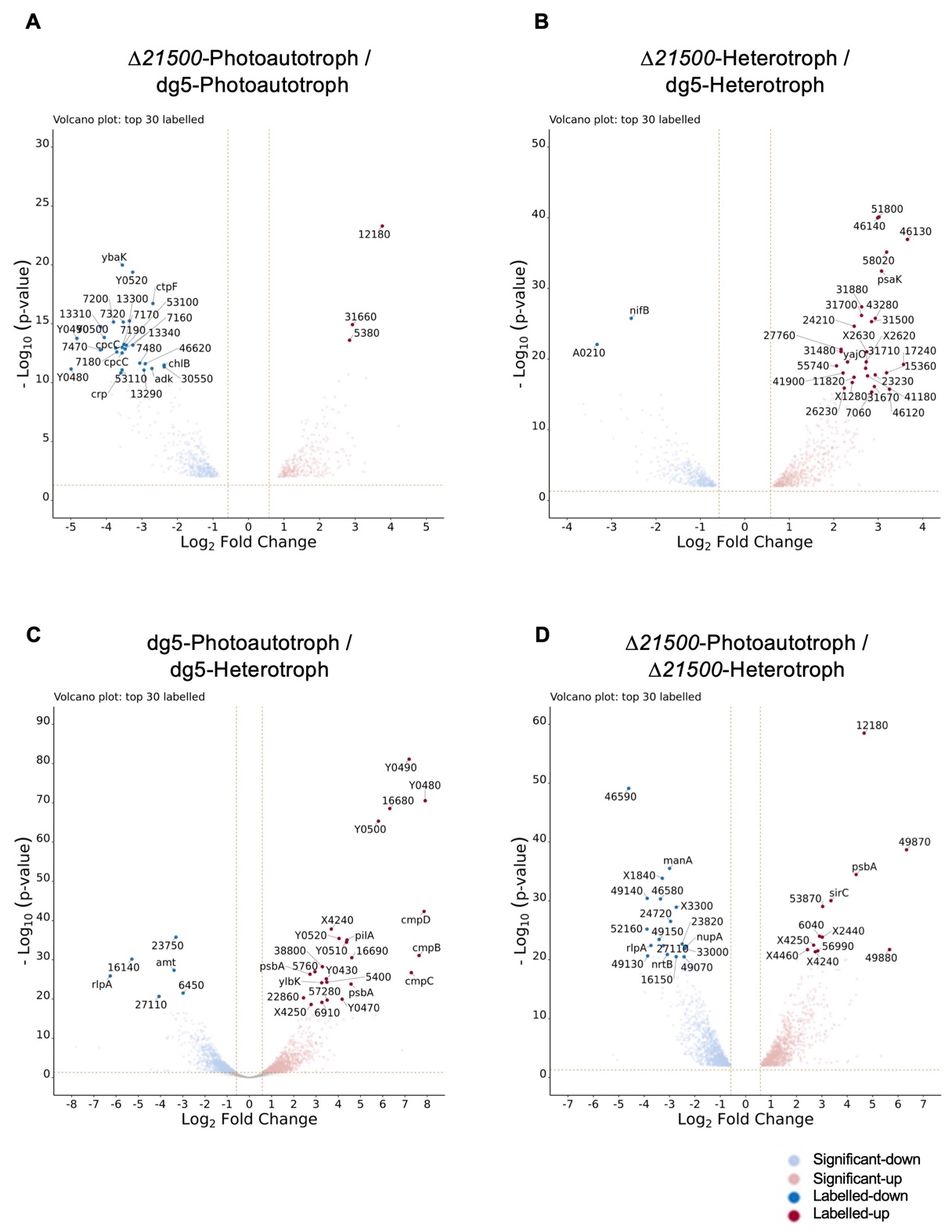

**Figure S7.**	Volcano plots of genes significantly differentially expressed by the knockout of *LBDG_21500* under light and dark conditions and in response to light and dark conditions of dg5 and *∆21500*. DEGs with higher and lower transcript levels in *∆21500* are shown in red and blue, respectively, compared with dg5 under photoautotrophic (**A**) and with dark-heterotrophic (**B**) conditions. In dg5 (**C**) and *∆21500* (**D**), DEGs with higher transcript levels in *∆21500* are shown in red and those with lower transcript levels in *∆21500* are shown in blue. Panels A and B are identical to those in Figure 6B, a and b, respectively. Volcano plots were created as described in Figure 6B.

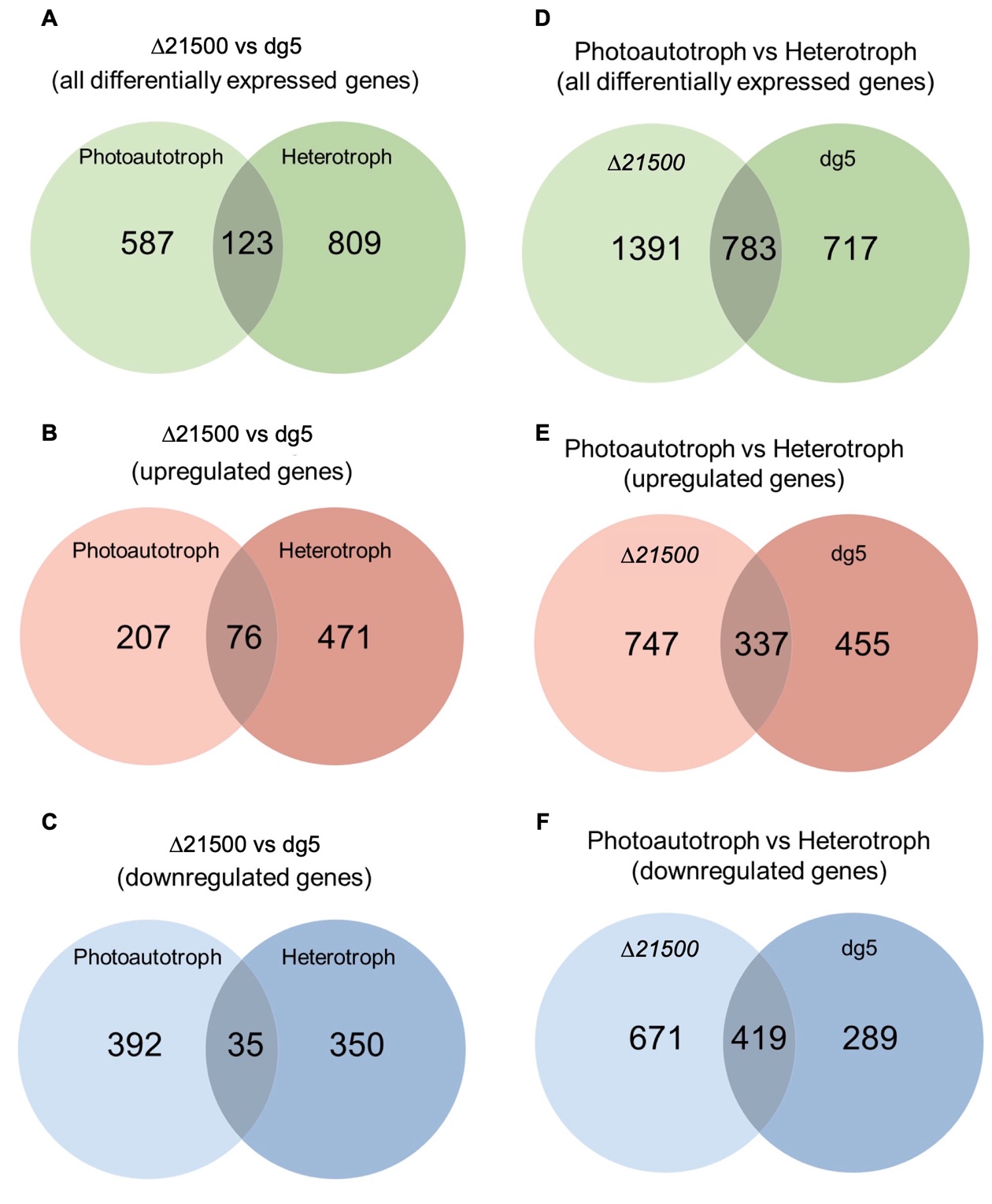

**Figure S8.**	Venn diagrams showing the number DEGs under each condition. DEG analysis was performed at a significance level of FDR < 0.01. (**A–C).** The total number of DEGs (**A**), the number of DEGs with higher transcript levels in *∆21500* relative to dg5 (**B**), and the number of DEGs with lower transcript levels in *∆21500* relative to dg5 (**C**) under photoautotrophic (left) and dark-heterotrophic (right) conditions. (**D–F).** The total number of DEGs (**D**), the number of DEGs with higher transcript levels under photoautotrophic conditions relative to dark-heterotrophic conditions (**E**), and the number of DEGs with lower transcript levels under photoautotrophic conditions relative to dark-heterotrophic (**C**) conditions in *∆21500* (left) and dg5 (right).

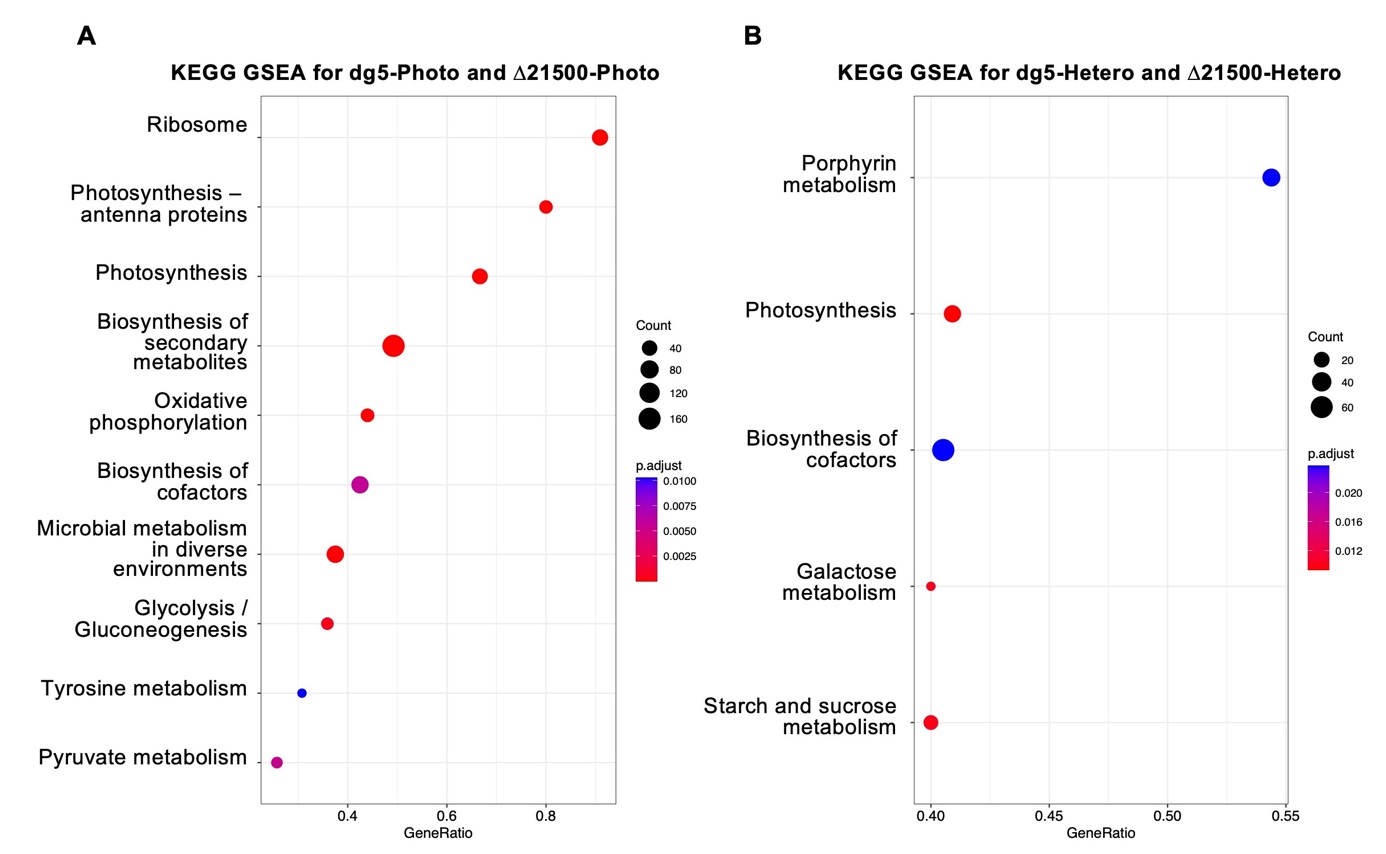

**Figure S9.**	Results of gene set enrichment analysis (GSEA). KEGG pathways that varied significantly under light (**A**) and dark (**B**) conditions were examined using preranked GSEA at a significance level of FDR < 0.01. The enriched pathway was plotted as a dot using the clusterProfiler package. The size of the plot shows the number of genes in the pathway. The plots are colored in gradations according to the FDR.

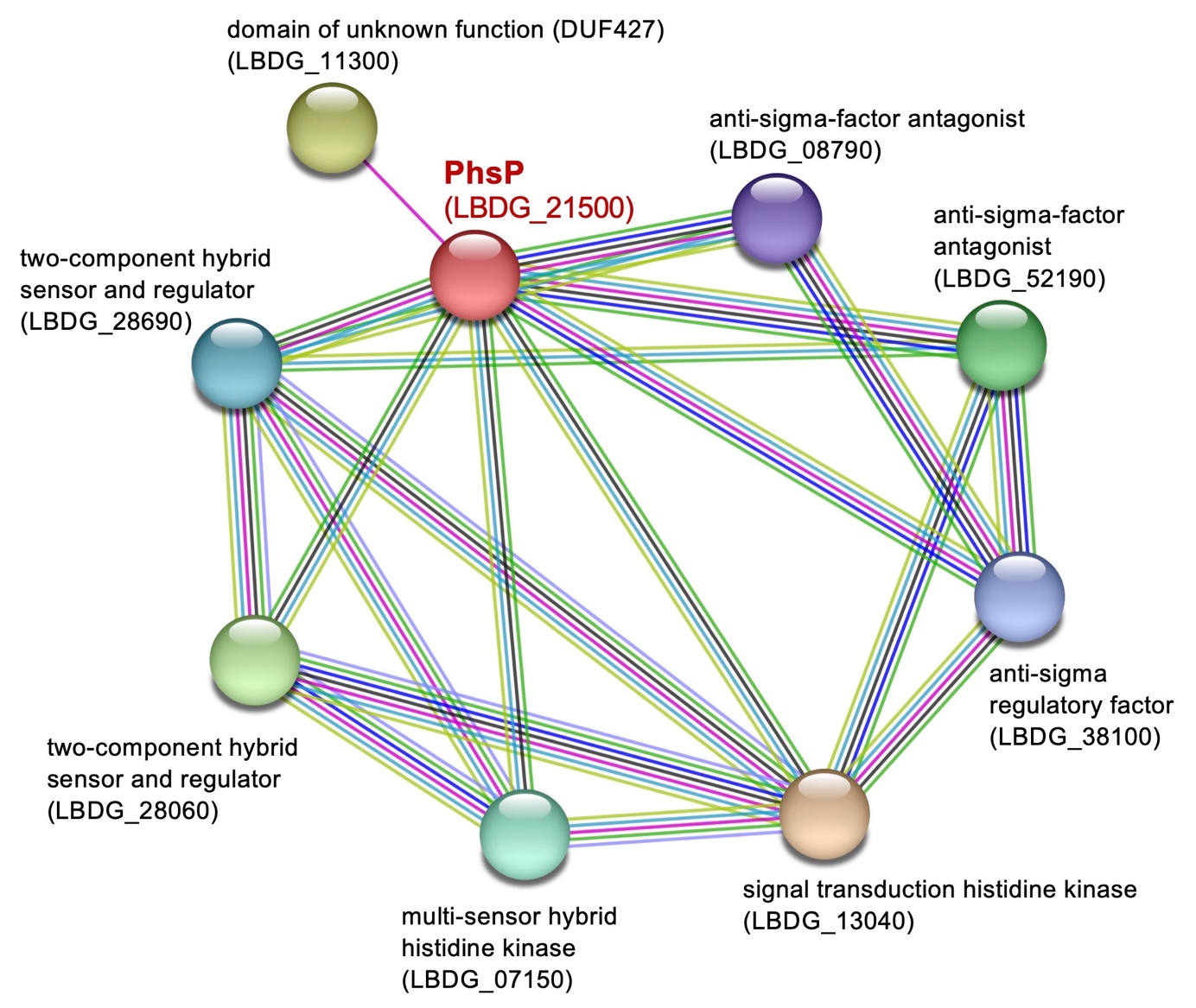

**Figure S10.** Protein–protein interaction (PPI) network analysis of PhsP-related proteins. Proteins that may interact with PhsP were explored using the STRING web interface (Szklarczyk, 2021; accessed in June 2022). PPI network analysis using STRING was used to search for proteins with which RsbU interacts. The PPI network of *L. boryana* PCC 6306, the closest relative species to *L. boryana*, was used for the analysis. The network shows 20 interactions formed by 9 proteins. Each interaction determined based on database (light blue), experimental results (purple), adjacent genes (green), co-occurrence (blue), text mining (yellow-green), and co-expression (black) corresponds to the color of the line connecting the proteins. RsbU proteins are circled in red, and other proteins are indicated by name and corresponding gene ID.
