## Supplementary Tables S1 and S2 for "Microevolution toward loss of photosynthesis: Mutations promoting dark-heterotrophic growth and suppressing photosynthetic growth in cyanobacteria"

**Table S1.** List of mutations in genomes of v1-v6.

| Predicted mutations | | | | | | | | | | | | | |
|---|---|---|---|---|---|---|---|---|---|---|---|---|---|
| seq id | position | mutation | WT | dg5 | v1 | v2 | v3 | v4 | v5 | v6 | annotation | gene | description |
| chr | 156,667 | T→C | +/- | +/- | +/- | +/- | +/- | + | +/- | +/- | intergenic (-330/+409) | LBWT_1250 ← / ← LBWT_1260 | hypothetical protein/hypothetical protein |
| | 1,305,274 | A→G | | + | + | + | + | + | + | + | V15A (GTC→GCC) | LBWT_11960 ← | hypothetical protein |
| | 1,535,169 | (A)8→6 | + | + | + | + | + | + | + | + | intergenic (-85/+149) | LBWT_14410 ← / ← LBWT_14420 | acyl-ACP reductase/fatty aldehyde decarbonylase |
| | 1,753,599-1,753,609 | Tn element | | | | + | | | | | intergenic(+81/+149) | LBWT_16400 → /LBWT_16410 → | carbamoyl-phosphate synthase small subunit/anti-sigma-factor antagonist |
| | 1,886,569 | C→T | + | | | | | | | | Y61Y (TAC→TAT) | LBWT_17680 → | hypothetical protein |
| | 1,952,058 | A→G | | | | | | | | + | T28A (ACC→GCC) | LBWT_18240 → | 30S ribosomal protein S11 |
| | 2,192,583 | +A | | | + | | | | | | coding (706/3276 nt) | LBWT_20410 → | Phycobilisome protein,Phycobilisome Linker polypeptide |
| | 2,318,951 | T→C | | | | | | + | | | D407G (GAC→GGC) | LBWT_21500 ← | serine phosphatase RsbU, regulator of sigma subunit |
| | 2,319,287 | C→T | | | + | | | | | | G295D (GGT→GAT) | | |
| | 2,319,683 | (TTTCTCA)1→2 | | | | | | | + | | coding (488/1425 nt) | | |
| | 2,320,114-2,320,120 | Tn element | | | | | + | | | | frameshift | | |
| seq id | position | mutation | WT | dg5 | v1 | v2 | v3 | v4 | v5 | v6 | annotation | gene | description |
| chr | 2,431,026 | (T)7→8 | + | + | + | + | + | + | + | + | intergenic (+62/+20) | LBWT_22630 → / ← LBWT_22640 | glutathione S-transferase/Photosystem II 4 kDa reaction center component superfamily |
| | 2,551,182 | Tn element | | | | | + | | | | intergenic (-53/-92) | LBWT_23800 ← / → LBWT_23810 | lysophospholipase L1-like esterase/DNA-directed RNA polymerase, subunit K/omega |
| | 2,565,721 | C→T | | + | + | + | + | + | + | + | R557K (AGA→AAA) | LBWT_23950 ← | PAS domain S-box |
| | 2,614,187 | Tn element | | | | | | | | + | intergenic (-66/+88) | LBWT_24380 ← / ← LBWT_24390 | Protein of unknown function (DUF2973)/hypothetical protein |
| | 2,628,331 | A→C | | | | | + | | | | intergenic (-83/-305) | LBWT_24480 ← / → LBWT_24490 | two component LuxR family transcriptional regulator/hypothetical protein |
| | 2,630,562 | (A)7→8 | + | + | + | + | + | + | + | + | intergenic (+7/-137) | LBWT_24500 → / → LBWT_24510 | adenosylcobyric acid synthase (glutamine-hydrolysing)/hypothetical protein |
| | 2,754,184 | A→G | + | + | + | + | + | + | + | + | G76G (GGA→GGG) | LBWT_25700 → | GD21975 |
| | 2,829,307 | TCTC→GAGA | | | | + | + | | | | coding (58-61/621 nt) | LBWT_26360 ← | DnaJ, molecular chaperone |
| | 2,880,914 | Tn element | | | | | | + | | | intergenic (-210/-312) | LBWT_26870/LBWT_26880 | complementary adaptation response regulator homolog/phycobilin lyase, CpcS-I subunit |
| | 3,151,311 | A→G | | + | + | + | + | + | + | + | K148E (AAA→GAA) | LBWT_29570 → | acid phosphatase/vanadium-dependent haloperoxidase related protein |
| seq id | position | mutation | WT | dg5 | v1 | v2 | v3 | v4 | v5 | v6 | annotation | gene | description |
| chr | 3,346,480 | G→A | | | | | | | + | | noncoding (47/74 nt) | LBWT_31360 → | tRNA-Pro |
| | 3,393,956 | G→A | | | | | | | + | | intergenic (-147/-147) | LBWT_31810 ← / → LBWT_31820 | integrase family protein/hypothetical protein |
| | 3,405,613 | Δ1,012 bp | + | | | | | | | | deletion | LBWT_31910 – LBWT_31920 | ABC-type nitrate sulfonate bicarbonate transport system, periplasmic component/transposase, IS4 family protein |
| | 3,453,673 | T→C | | + | + | + | + | + | + | + | intergenic (-10/-266) | LBWT_32410 ← / → LBWT_32420 | folate/biopterin transporter/Chlorophyll A-B binding protein |
| | 3,486,871 | Δ1 bp | | + | | | | | | | coding (1967/1989 nt) | LBWT_32770 → | putative sensor protein |
| | 3,573,479-3,573,484 | Tn element | | | | + | | | | | frameshift | LBWT_33610 ← | putative P-loop ATPase |
| | 4,517,660-4,523,614 | Δ5,965 bp | | | | | | | | + | deletion | LBWT_42760–LBWT_42810 | trypsin-like serine protease with C-terminal PDZ domain/ peptidase membrane zinc metallopeptidase<br>metal dependent phosphohydrolase/ iojap-like ribosome-associated protein<br>hypothetical protein/ homodimeric phycobilin lyase, CpcS-III family |
| | 4,619,261 | Δ1 bp | | | | | | + | | | coding (715/762 nt) | LBWT_43870 ← | Phycobilisome Linker polypeptide |
| | 5,175,200 | +T | | + | + | + | + | + | + | + | coding (46/294 nt) | LBWT_49050 ← | cytochrome c, class I, cytochrome cM |
| | 5,325,830 | G→A | | | | | | + | | | E190K (GAG→AAG) | LBWT_50360 → | CheA Signal Transduction Histidine Kinases (STHK) |
| seq id | position | mutation | WT | dg5 | v1 | v2 | v3 | v4 | v5 | v6 | annotation | gene | description |
| chr | 6,085,873-6,085,876 | Tn element | | | | | | | | + | frameshift | LBWT_57340 → | bacteriophytochrome (light-regulated signal transduction histidine kinase) |
| pLBY | 44,358 | A→G | | | | | | + | | | intergenic (-84/–) | LBWT_Y0550 ← / – | hypothetical protein/– |
| pLBY | 44,362 | G→A | | | | | | + | | | intergenic (-88/–) | LBWT_Y0550 ← / – | hypothetical protein/– |

**Table S2.** List of mutations in genomes of dg201-dg229.

Predicted mutations

| seq id | position | mutation | WT | 201 | 202 | 204 | 205 | 206 | 207 | 208 | 210 | 211 | 214 | 217 | 218 | 219 | 220 | 222 | 223 | 224 | 225 | 226 | 227 | 228 | 229 | effect | gene | description |
|---|---|---|---|---|---|---|---|---|---|---|---|---|---|---|---|---|---|---|---|---|---|---|---|---|---|---|---|---|
| chr | 244,718 | A→G | | | | + | | | | | | | | | | | | | | | | | | | | intergenic (-156/-163) | LBWT_2100 ← / → LBWT_2110 | glycyl-tRNA synthetase alpha chain/16S ribosomal RNA |
| chr | 244,731 | 2 bp→TC | | | | + | | | | | | | | | | | | | | | | | | | | intergenic (-169/-149) | | |
| chr | 433,335 | T→A | | | | + | | | | | | | | | | | | | | | | | | | | N884Y (AAC→TAC) | LBWT_4010 ← | putative low-complexity protein |
| chr | 557,633 | C→T | | | | | | | | | | | | | | + | | | | | | | | | | P67S (CCC→TCC) | LBWT_5210 → | Cytochrome c biogenesis protein ccsA |
| chr | 557,674 | G→A | | | | | | | | + | | | + | | | | | | | | | | | | | W80* (TGG→TGA) | | |
| chr | 557,820 | C→T | | | | | | | | | | | | | | | + | | | | | | | | | P129L (CCT→CTT) | | |
| chr | 557,892 | G→A | | | | | | | | | | | | | + | | | | | | | | | | | G153E (GGA→GAA) | | |
| chr | 558,170 | G→A | | | | + | | | | | | | | | | | | | | | | | | | | A246T (GCG→ACG) | | |
| chr | 677,198 | G→A | | | | | | | + | | | | | | | | | | | | | | | | | intergenic (+144/+178) | LBWT_6320 → / ← LBWT_6330 | hypothetical protein/transposase, IS4 family |
| chr | 677,200 | T→A | | | | | | | + | | | | | | | | | | | | | | | | | intergenic (+146/+176) | | |
| chr | 677,202 | T→C | | | | | | | + | | | | | | | | | | | | | | | | | intergenic (+148/+174) | | |
| chr | 808,530 | Tn element | | | | | | | | | + | | | | | | | | | | | | | | | coding (202/831 nt) | LBWT_7420 ← | oxidoreductase FAD/NAD(P)-binding domain-containing protein |
| chr | 899,137 | A→G | | | | | | | | | | | | | | | | | + | | | | +/- | | | R78R (CGA→CGG) | LBWT_8190 → | hypothetical protein |
| chr | 910,863 | T→A | | | + | | | | | | | | | | | | | | + | | | | | | | intergenic (-345/+19) | LBWT_8330 ← / ← LBWT_8350 | hypothetical protein/hypothetical protein |
| chr | 1,319,558 | C→G | | | | | | | + | | | | | | | | | | | | | | | | | S76S (TCC→TCG) | LBWT_12200 → | helix-turn-helix domain-containing protein |
| chr | 1,350,776 | T→A | | | + | | | | | | | | | | | | | | + | | | | | | | I1900F (ATC→TTC) | LBWT_12620 ← | TP901 family prophage L54a |
| chr | 1,442,767 | C→T | | | +/- | | | | | | | | | | | | | | + | | | | | | | W284* (TGG→TGA) | LBWT_13590 ← | Cytochrome c biosynthesis protein CcsB |
| chr | 1,442,933 | A→T | | | +/- | | | | | | | | | | | | | + | | | | | | | | V229E (GTG→GAG) | | |
| chr | 1,443,605 | Tn element | | | | | | | + | | | | | | | | | | | | | | | | | coding (14/1371 nt) | | |
| chr | 1,535,169 | (A)$^{8\to6}$ | + | + | + | + | + | + | + | + | + | + | + | + | + | + | + | + | + | + | + | + | + | + | + | intergenic (-85/+149) | LBWT_14410 ← / ← LBWT_14420 | acyl-ACP reductase/fatty aldehyde decarbonylase |
| seq id | position | mutation | WT | 201 | 202 | 204 | 205 | 206 | 207 | 208 | 210 | 211 | 214 | 217 | 218 | 219 | 220 | 222 | 223 | 224 | 225 | 226 | 227 | 228 | 229 | effect | gene | description |
| chr | 1,621,764 | 18 bp IS | | | | + | | | | | | | | | | | | | | | | | | | | coding (569/2052 nt) | LBWT_15240 ← | histidine kinase with GAF domain |
| chr | 1,753,741 | +C | | | | | | | | | | | | | | + | | | | | | | | | | coding (15/252 nt) | LBWT_16410 → | anti-sigma-factor antagonist |
| chr | 1,886,569 | C→T | + | | | | | | | | | | | | | | | | | | | | | | | Y61Y (TAC→TAT) | LBWT_17680 → | hypothetical protein |
| chr | 2,243,093 | (GCCATC)$^{2\to1}$ | | + | | | | | | | | | | | | | | | | | | | | | | coding (832-837/1635 nt) | LBWT_20850 ← | phosphoglucomutase |
| chr | 2,317,726-2,318,922 | Δ1197 bp | | | | | | | | | | | | | | | | | | + | | | | | | coding (948/951 nt)-(1248/1425 nt) | LBWT_21480-LBWT_21490-LBWT_21500 | glycosyl transferase/transcriptional regulator, GntR family/serine phosphatase RsbU |
| chr | 2318614-2,319,607 | Δ993 bp | | | | | | | | | | | | | | | | | | | | | | + | | coding(601/672 nt)-(564/1425 nt) | LBWT_21490 →/←LBWT_21500 | transcriptional regulator, GntR family/serine phosphatase RsbU |
| chr | 2,318,853 | G→A | | | | | | | + | | | | | | | | | | | | | | | | | Q440* (CAG→TAG) | LBWT_21500 ← | serine phosphatase RsbU, regulator of sigma subunit |
| chr | 2,319,064 | Δ2 bp | | | | | | | | | | | + | | | | | | | | | | | | | coding (1106-1107/1425 nt) | | |
| chr | 2,319,028 | Tn element | | | | | | | | + | | | | | | | | | | | | | | | | coding (1143/1425 nt) | | |
| chr | 2,319,079 | C→T | | | | | | + | | | | | | + | | | | | | | | | | | | W364* (TGG→TGA) | | |
| chr | 2,319,099 | G→A | | | | | | | | | | | | | | | + | | | | | | | | | H358Y (CAC→TAC) | | |
| chr | 2,319,149-2,319,161 | Tn element | | | | | | | | | | | | | | | | + | | | | | | | | coding (1016-1022/1425 nt) | | |
| chr | 2,319,227 | G→A | | | | | | | | | | + | | | | | | | | | | | | | | P315L (CCC→CTC) | | |
| chr | 2,319,305 | C→T | | | | | + | | | | | | | | | | | | | | | | | | | G289D (GGC→GAC) | | |
| chr | 2,319,337 | +A | | | | | | | | | | | | | | | | | | | + | | | | | coding (834/1425 nt) | | |
| chr | 2,319,419-2,319,427 | Tn element | | | | | | | | | | | | | | | | | | | | | + | | | coding (744-752/1425 nt) | | |
| chr | 2,319,515 | Tn element | | | | | | | | | | | | | | | | | | | | + | | | | coding (656/1425 nt) | | |
| chr | 2,319,958 | Δ1 bp | | | | | | | | | | | | | + | | | | | | | | | | | coding (213/1425 nt) | | |
| chr | 2,360,415 | 3 bp→TCA | | | | | | | + | | | | | | | | | | | | | | | | | intergenic (-32/+232) | LBWT_21910 ← / ← LBWT_21920 | 16S ribosomal RNA/1-acyl-sn-glycerol-3-phosphate acyltransferase |
| chr | 2,360,419 | A→T | | | | | | | + | | | | | | | | | | | | | | | | | intergenic (-36/+230) | | |
| chr | 2,360,421 | Δ1 bp | | | | | | | + | | | | | | | | | | | | | | | | | intergenic (-38/+228) | | |
| seq id | position | mutation | WT | 201 | 202 | 204 | 205 | 206 | 207 | 208 | 210 | 211 | 214 | 217 | 218 | 219 | 220 | 222 | 223 | 224 | 225 | 226 | 227 | 228 | 229 | effect | gene | description |
| chr | 2,381,928-2,381,936 | Tn element | | | | | | | | | | | | | | | + | | | | | | | | | coding (1466-1474/1959 nt) | LBWT_22130 ← | PAS domain-containing protein |
| chr | 2,424,616 | C→T | | | | | | | | + | | + | | | | | | | | | | + | + | + | | P98P (CCG→CCA) | LBWT_22570 ← | C-3',4' desaturase CrtD |
| chr | 2,431,026 | (T)$^{7\to8}$ | + | + | + | + | + | + | + | + | + | + | + | + | + | + | + | + | + | + | + | + | + | + | + | intergenic (+62/+20) | LBWT_22630 → / ← LBWT_22640 | glutathione S-transferase/Photosystem II 4 kDa reaction center component superfamily |
| chr | 2,566,519 | G→A | | + | | | | | | | | | | | | | | | | | | | | | | A291V (GCC→GTC) | LBWT_23950 ← | PAS domain S-box |
| chr | 2,604,965 | (ATCG)$^{2\to3}$ | | + | + | + | + | + | + | + | + | + | + | + | + | + | + | + | + | + | + | + | + | + | + | coding (1100/1269 nt) | LBWT_24300 ← | glycosyl transferase group 1 |
| chr | 2,630,562 | (A)$^{7\to8}$ | + | + | + | + | + | + | + | + | + | + | + | + | + | + | + | + | + | + | + | + | + | + | + | intergenic (+7/-137) | LBWT_24500 → / → LBWT_24510 | adenosylcobyric acid synthase (glutamine-hydrolysing)/hypothetical protein |
| chr | 2,634,859 | C→T | | | | | | | | | | | | | | | | | + | | | | | | | G169D (GGC→GAC) | LBWT_24540 ← | arabinose efflux permease family protein |
| chr | 2,754,184 | A→G | + | + | + | + | + | + | + | + | + | + | + | + | + | + | + | + | + | + | + | + | + | + | + | G76G (GGA→GGG) | LBWT_25700 → | GD21975 |
| chr | 3,345,295 | G→C | | | | | | | | | | | | | | | | | + | | | | | | | M530I (ATG→ATC) | LBWT_31350 → | DNA topoisomerase I, bacterial |
| chr | 3,405,613 | Δ1,012 bp | + | + | + | + | + | + | + | + | + | + | + | + | + | + | + | + | + | + | + | + | + | + | + | coding(1276/1293 nt)-intergenic | [LBWT_31910] → / → LBWT_31920 | ABC-type nitrate/sulfonate/bicarbonate transport system/transposase, IS4 family protein |
| chr | 3,531,743 | T→G | | | | | | | | + | | + | | | | | | | | | | + | + | + | | D120A (GAC→GCC) | LBWT_33180 ← | response regulator containing CheY-like receiver domain and HTH DNA-binding domain |
| chr | 3,861,797 | Tn element | | | | | | | | | | + | | | | | | | | | | | | | | coding (27/1503 nt) | LBWT_36280 → | carotene isomerase CrtH |
| chr | 4,699,723 | G→A | | | | | | + | | | | | | | | | | | | | | | | | | R5C (CGC→TGC)<br>S945L (TCG→TTG) | LBWT_44580 ←<br>LBWT_44590 ← | Adenylate and Guanylate cyclase catalytic domain protein<br>multi-sensor hybrid histidine kinase |
| chr | 4,994,433 | C→T | | | | | + | | | | | | | | | | | | | | | | | | | T95I (ACA→ATA) | LBWT_47500 → | prepilin-type N-terminal cleavage/methylation domain-containing protein |
| chr | 5,175,285 | Tn element | | | | | | | | | + | | | | | | | | | | | | | | | intergenic(+40/+489) | LBWT_49050 ←/→LBWT_49055 | cytochrome c, class I, cytochrome cM/Cytochrome b6-f complex subunit 5, PetG |
| chr | 5,634,330 | +A | | | | | | | | | | | | + | | | | | | | | | | | | coding (881/1416 nt) | LBWT_53250 → | phytoene desaturase CrtP |
| chr | 5,634,938 | +A | | | | | + | | | | | | | | | | | | | | | | | | | coding (29/936 nt) | LBWT_53260 → | phytoene synthase CrtB |
| pLBX | 337,028 | Δ1 bp | | | | | | | | | | | | + | | + | | | | | + | | | | | coding (2220/6021 nt) | LBWT_X2810 ← | Serine/Threonine protein kinase and Signal Transduction Histidine Kinase |
| seq id | position | mutation | WT | 201 | 202 | 204 | 205 | 206 | 207 | 208 | 210 | 211 | 214 | 217 | 218 | 219 | 220 | 222 | 223 | 224 | 225 | 226 | 227 | 228 | 229 | effect | gene | description |
| pLBY | 44,358 | A→G | | + | | | | | | | | | | | | | | | | | | | | | | intergenic (-84/–) | LBWT_Y0550 ← / – | hypothetical protein/– |
| pLBY | 44,362 | G→A | | | | | | | | | + | | | | | | | | | | | | | | | intergenic (-88/–) | | |
